# Reduced legacy precipitation decreases microbial community growth efficiency and alters soil organic carbon in a California grassland

**DOI:** 10.1101/2025.09.23.678135

**Authors:** Linnea K. Hernandez, Nicole Didonato, Ljiljana Pasa-Tolic, Pete F. Chuckran, Mary K. Firestone, Ella T. Sieradzki, Mengting Maggie Yuan, Katerina Estera-Molina, Jeffrey Kimbrel, Paul Dijkstra, Jillian F. Banfield, Jennifer Pett-Ridge, Steven J. Blazewicz

**Affiliations:** Physical and Life Science Directorate, Lawrence Livermore National Laboratory, Livermore, CA, USA; Environmental Molecular Science Laboratory (EMSL), Earth and Biological Sciences Division, Pacific Northwest National Laboratory, Richland, WA, USA; Department of Environmental Science, Policy, and Management, University of California, Berkeley, CA, USA; Earth and Environmental Sciences, Lawrence Berkeley National Laboratory, Berkeley, CA, USA; Department of Agroecology, Aarhus University, Aarhus, Denmark; Pacific Biosciences Research Center, University of Hawaii, Manoa, HI, USA; Center for Ecosystem Science and Society (ECOSS) and Department of Biological Sciences, Northern Arizona, University, Flagstaff, AZ, USA; Life & Environmental Science Department, University of California Merced, Merced, CA, USA; Innovative Genomics Institute, University of California, Berkeley, Berkeley, CA, USA

## Abstract

Changes in global patterns can leave a lasting legacy in semi-arid grasslands by reshaping microbial growth dynamics and carbon cycling during the first wet-up in the autumn—a period known for intense microbial activity and significant carbon emissions. To study the lasting impacts of decreased winter rain, we implemented two precipitation regimes (100% vs. 50% mean annual precipitation) in California Mediterranean-climate grassland field plots. After the dry season, soils were rewetted in the laboratory with H_2_^18^O, and sampled at 0 h, 3 h, 24 h, 48 h, 72 h, and 168 h post rewet. We quantified CO_2_ efflux; measured microbial growth and mortality via quantitative ^18^O stable isotope probing and 16S rRNA gene amplicon sequencing; and characterized the soil organic carbon chemical composition, metagenomes, and metatranscriptomes. We found that reduced winter precipitation imposed a strong legacy effect on microbial turnover; despite maintaining similar respiration rates, microbial growth declined by ∼1 order of magnitude, yielding decreased community growth efficiency (CGE = gross community growth/net respiration), and microbial mortality declined by ∼2 orders of magnitude. Soil organic carbon also shifted from lipid-like, amino-sugar-like, and protein-like compounds (indicative of microbial necromass) to more oxidized lignin-like and tannin-like compounds (indicative of decomposing plant-derived compounds). Meta-omics revealed distinct metabolic strategies linked to CGE. At high-CGE, microbes appeared to consume more energetically favorable N-rich necromass (released via high microbial turnover), this allowed for increased amino acids and peptidoglycan biosynthesis and greater aromatic compound degradation, fueling further energy production and growth efficiency. At low-CGE, communities had elevated carbohydrate metabolism and lipid turnover, consistent with increased investment in plant detritus degradation and membrane repair and maintenance rather than growth. Together, our findings demonstrate that reduced winter rainfall decreases microbial turnover following rewetting. Persistent decreases in CGE due to reduced winter rainfall result in consistent carbon loss as CO_2_, which, if sustained over multiple years, could ultimately lead to a net decline in total soil organic carbon.

## 1. Introduction

Climate change is projected to alter global precipitation patterns, leading to more frequent droughts and more intense rewetting events [1]. These changes may have profound implications for soil microbial communities which are central to belowground biogeochemical cycling [2, 3]. Microbial diversity and activity are highly sensitive to moisture conditions [4, 5], and deviations from typical moisture regimes are likely to influence carbon cycling and soil organic carbon (SOC) storage [6].

Microbes control soil carbon fate through partitioning of carbon between anabolism (biosynthesis of biomass for growth and secondary metabolites), and catabolism (molecule breakdown and production of CO_2_ via energy-generating pathways). In soils, microbial growth followed by mortality leads to the formation of microbial necromass which can account for up to 50% of total SOC [3, 7], highlighting the importance of microbial turnover in SOC accumulation and persistence. One quantitative measure of microbial C allocation to growth versus respiration is carbon use efficiency (CUE; C-biomass ÷ [C-biomass + C-CO_2_]). Several studies have found that higher CUE is positively correlated with overall SOC [8, 9] indicating that increased C allocation to biomass enhances SOC. However, this relationship is not consistent; for example, mineral-associated organic carbon has been found to be both positively [10] and negatively [11] associated with CUE. These discrepancies may reflect different CUE measurement approaches [12], but could also be due to differences in substrate, enzyme investment, and possible energy-spilling reactions.

Soil moisture regulates microbial growth and respiration because water governs nutrient diffusion, chemical reactions, and cellular integrity. Under drought conditions, microbial community composition [13, 14] and gene expression [6, 15] often shift, with concurrent decreases in growth and respiration [16] which could leave CUE unchanged, as found in some grassland ecosystems [17, 18]. In contrast, studies in forest, agricultural, and some grassland ecosystems have reported increased CUE under drought, which is attributed to reduced respiration yet unchanged growth rates [18, 19]. These findings indicate that the effects of drought on microbial activity and CUE may be ecosystem dependent and could differ based on timing and length of droughts.

Beyond drought, seasonal fluctuations in soil moisture can further affect the dynamics of microbial growth and respiration. Seasonal changes in soil moisture, with alternating periods of dry and wet conditions, exert strong controls on microbial processes. For example, in grassland ecosystems, drier summer conditions often lead to reduced microbial growth compared with wetter winter conditions [4, 19]. While some seasonal shifts in soil moisture occur gradually, abrupt shifts, such as rewetting after a dry period, can have dramatic effects on microbial activity and carbon dynamics. Rewetting events often trigger a burst of microbial respiration, frequently referred to as the “Birch effect” [20]. This results in substantial soil CO_2_ efflux, equivalent to as much as 10% of annual net ecosystem production [21, 22]. In a California grassland, measurements made after rewetting showed a simultaneous burst in microbial growth and mortality, indicating high microbial turnover [23, 24]. These findings highlight the potential for rewetting to significantly influence both C release to the atmosphere and the generation of microbial necromass—a key precursor for persistent soil carbon.

While the immediate effects of soil moisture on microbial growth and respiration have been well studied, the legacy effects of moisture regimes—how past moisture conditions influence current microbial activity—remain less understood. The concept of “ecological memory” describes how past events influence current ecosystem responses; this concept is prevalent in research on aboveground ecosystems such as plants [25] and is increasingly applied to soil microbiomes [26–29]. Canarini et al. [26] reported that the microbial community composition differed between soils subjected to the 1^st^ drought and those subjected to the 10^th^ drought, leading to altered predicted functional capacity and enzyme production. However, other studies, such as Liu et al. [30], reported no differences in microbial composition or respiration between legacy drought and control soils after one month, indicating variability in legacy effects. More studies are needed to better understand how these precipitation legacy effects influence microbial turnover and microbial carbon allocation at the field-scale.

An important step in this direction is to improve our mechanistic understanding of the drivers of CUE and microbial carbon allocation. We expect CUE to be tightly regulated by i) substrate properties including availability, thermodynamics, and elemental stoichiometry; ii) current and legacy moisture; and iii) how microbial communities allocate transcription toward biosynthesis, energy generation, and enzyme production. With regard to substrate properties, pure-cultures and mono-substrate incubations showed that glucose yielded higher CUE, whereas an aromatic carbon source yielded lower CUE [31]. Other work has shown that substrates with higher C:N suppress CUE because microbes respire excess carbon to obtain nitrogen [32]. Yet, studies that simultaneously measure SOC chemical composition and active microbial pathways alongside community-scale growth efficiency in soils are lacking.

Furthermore, it is critical to employ measurement approaches that capture the full scope of microbial carbon processing across the whole community under natural soil conditions. To address this, our study adopts the term community growth efficiency (CGE) to describe the ratio of newly synthesized microbial biomass, measured with H_2_^18^O DNA quantitative stable isotope probing (qSIP), to total CO_2_ released, which represents carbon mineralized for energy to support growth and cell maintenance [33]. Unlike traditional CUE measurements, which typically rely on single substrate incubations (i.e. ^13^C-glucose) and aim to quantify the absolute efficiency of carbon conversion [33], CGE integrates carbon uptake from all available sources in situ and reflects the community-level allocation of carbon under natural soil conditions. While the absolute values of CGE are not directly comparable to those reported for CUE in the literature, the directional trends in CGE provide meaningful insights into how microbial communities respond to environmental changes, such as drought and rewetting, in terms of carbon allocation. By comparing CGE trends with established CUE patterns, we aim to contextualize our findings within the broader framework of microbial carbon cycling in soils, while maintaining transparency about the methodological distinctions inherent to our approach.

In this study, we investigated the legacy effects of normal and reduced precipitation regimes during the winter growing season on microbial growth dynamics following rewetting after the summer dry period in a California annual grassland with a Mediterranean-type climate. Specifically, we examined how these past precipitation regimes influenced (1) taxon-specific microbial growth and mortality via 16S rRNA gene quantitative stable isotope probing (qSIP), (2) CGE (microbial net growth divided by total CO_2_ efflux), (3) SOC chemical composition, and (4) microbial gene expression via metatranscriptomics. We hypothesized that reduced precipitation during the winter growing season would have a legacy effect, decreasing microbial growth and mortality during rewetting six months later. Additionally, we hypothesized that these changes would influence SOC profiles and microbial activity; soils exposed to normal precipitation regimes would have higher CGE and exhibit increased expression of genes associated with biosynthesis, whereas soils exposed to reduced precipitation regimes would have lower CGE and increased expression of genes associated with energy acquisition. By improving our understanding of how reduced rainfall impacts microbial growth, carbon allocation, and SOC composition dynamics, this research provides critical insights into how climate change may affect SOC persistence. These findings can inform landscape management strategies aimed at increasing SOC storage in soils.

## 2. Methods

### 2.1 Field site description

This field site is at the Hopland Research and Extension Center (HREC) located in northern California (39° 00’ 14.6” N, 123° 05’ 09.1” W) on territory originally occupied by the indigenous Pomo Nation. This site experiences a Mediterranean climate with cool, wet winters and warm, dry summers where most of annual precipitation occurs between October and March and most plant growth occurs from late February to early April [34]. Experimental plots were located on soils of the Squawrock–Witherell complex, have a pH of 5.9, total organic C content of 15.1 mg g^-1^, and total N content of 1.4 mg g^-1^ [35].

In October 2017, sixteen 3.24 m^2^ plots dominated by *Avena barbata* (wild oat grass) were randomly divided into two treatments that received either full (100% mean annual precipitation [MAP]; 930 mm) or reduced (50% MAP; 465 mm) precipitation. Precipitation was controlled controlled with rainout shelters constructed over each plot and a plaster pond liner placed belowground to 1 m depth to isolate each plot and reduce soil moisture flow between plots. The final rainfall occurred in May 2018, followed by the summer dry-down. In August 2018, at the end of the summer dry down when the gravimetric soil moisture was 3%, topsoil samples (0–15 cm, approximately 4700 cm^3^) were collected from eight plots (n = 4 each for 50 and 100% MAP; total of 48 samples) with cleaned shovels (rinsed with Eliminase [DeconLabs, King of Prussia, PA] and water between samples), immediately placed into jumbo-sized Ziplock bags, and transferred to Lawrence Livermore National Laboratory where roots were removed and soils were homogenized by sieving (2 mm) to remove rocks and large plant debris in a greenhouse with a dry and hot atmosphere.

### 2.2 Laboratory ^18^O-H_2_O wet-up and sample collection

The samples in the laboratory were subjected to H_2_^18^O labeling for quantitative growth analysis (for details, see [35–38]). Briefly, 5 g of sieved soil from each of the eight plots was transferred each to 11 Nalgene flat bottom vials (15 mL), for a total of 88 microcosms. One mL of isotopically enriched water (98-atom H_2_^18^O) or natural abundance water (control) was slowly and evenly pipetted onto the soil and gently mixed with the pipette tip, resulting in a final average gravimetric soil moisture of 22%. Two vials each were then immediately sealed inside a 500 mL mason jar with lids fitted with septa (to facilitate headspace sampling) and incubated at room temperature in the dark. Parallel jars (total of four vials) were destructively harvested at multiple timepoints following rewetting (0 h [before labeling/rewetting], 3 h, 24 h, 48 h, 72 h, and 168 h). For each harvest, vials containing soil were flash frozen in liquid nitrogen and stored at −80 °C.

Headspace gas samples (5 mL) were collected at each soil harvesting time point with a gas-tight syringe and transferred to 20 mL Wheaton serum bottles (DWK Life Sciences, Wertheim, Germany) that had been purged and filled with N_2_ (1 atm). Total headspace CO_2_ was quantified via gas chromatography equipped with a methanizer paired with a flame ionization detector (GC2015, Shimadzu).

### 2.3 16S-qSIP and metagenomics

#### 2.3.1 DNA extraction and fractionation

DNA was extracted from the soil samples via a modified phenol–chloroform protocol adapted from [39]. Each sample was extracted in triplicate, and the final DNA extracts were combined. For each extraction, soil (0.4 g) was added to a 2 mL Lysing Matrix E tube (MP Biomedicals, Irvine, CA) and extracted twice as follows: each tube received 500 µL extraction buffer (5% cetyltrimethylammonium bromide [CTAB], 0.5 M NaCl, 240 mM K_2_HPO_4_, pH 8.0) and 500 µL of 25:24:1 phenol:chloroform:isoamyl alcohol, shaken (30 s, 5.5 ms^-1^) (FastPrep24, MP Biomedicals, Santa Clara, CA), and then centrifuged for 5 min (16,100 × g). To remove residual phenol, the supernatant was added to a pre-spun 2 mL phase lock gel tube (5 prime, Gaithersburg, MD) with an equal volume of 24:1 chloroform:isoamyl alcohol, mixed by inverting, and centrifuged for 2 min (16,100 × g). The aqueous phases from both extractions were pooled, mixed with 7 µL of RNAse (10 mg/mL) by inverting, incubated for 1 h (4 °C), and centrifuged for 15 min (16,100 × g). The supernatant was added to a new 1.7 mL microcentrifuge tube, and 1 µL of glycoblue (15 mg/mL) and 1 mL of 40% polyethylene glycol (PEG) 6000 in 1.6 M NaCl were added, mixed by vortexing, and incubated at room temperature in the dark for 2 h. After centrifugation for 20 min (16,100 × g), the pellet was rinsed with 1 mL of ice-cold 70% ethanol, air-dried, resuspended in 30 µL of 1x Tris-EDTA (TE), and stored at −80 °C.

To measure the degree of ^18^O incorporation in the DNA, each sample was separated in a cesium chloride density gradient via ultracentrifugation, as previously described [39]. For each sample, 5 µg of DNA in 150 µL of 1xTE was mixed with 1 mL gradient buffer (0.1 mol L^-1^ Tris, 0.1 mol L^-1^ KCl, and 1 mmol L^-1^ EDTA) and 4.6 mL cesium chloride (CsCl) stock (1.885 g mL^-1^), with a final average density of 1.73 g mL^-1^. The samples (∼5.2 mL) were loaded into an ultracentrifuge tube (13 mm × 51 mm Quick-Seal, Beckman Coulter Brea, CA) and spun at 20 °C for 108 h at 176,284 RCF_avg_ in an Optima XE-90 ultracentrifuge (Beckman Coulter, Brea, CA) using a VTi65.2 rotor. Immediately following centrifugation, the contents of each tube were separated into 36 fractions (∼200 µL each) via a high-throughput, automated SIP pipeline (HT-SIP) [40]. Each tube was mounted in a fraction recovery system (Beckman Coulter, Brea, CA) where a 1260 isocratic pump (Agilent Technologies, Santa Clara, CA) delivered water at 0.25 mL min^-1^ through a 25G needle inserted through the top of the ultracentrifuge tube, and a side port needle routed to a 1260 Infinity (Agilent Technologies, Santa Clara, CA) fraction collector was inserted into the bottom of the tube to collect fractions in 96-well deep well plates. The density of each fraction was measured using an AR200 digital refractometer (Reichart, Depew, NY) fitted with a prism covering to facilitate measurement from 5 µL, as previously described [41]. The DNA in each fraction was purified and concentrated using a Hamilton Microlab Star liquid handling system (Hamilton Technology, Reno, NV) programmed to automate glycogen/PEG precipitation [40]. The washed DNA pellets were suspended in 40 µL of 1xTE, and the DNA concentration of each fraction was quantified via a PicoGreen fluorescence assay (Invitrogen, Waltham, MA). The fractions for each sample were pooled into five groups on the basis of density (1.6400-1.7039 g mL^-1^, 1.7040-1.7169 g mL^-1^, 1.7170-1.7299 g mL^-1^, 1.7300-1.7449 g mL^-1^, and 1.7450-1.7800 g mL^-1^), and the fractions within a pooled group were combined and sequenced.

#### 2.3.2 Quantitative PCR

Bacterial 16S rRNA gene copy abundances in the SIP fractions were quantified by qPCR in triplicate using the primers EUB 338F/EUB 518R [42] and averaged. Each 10 µL reaction contained 1X Forget-Me-Not EvaGreen qPCR Master Mix (Biotium, Fremont, CA), 1.5 mM MgCl_2_, 0.2 µM of each primer, and 1 µL of template. Reactions were performed on a CFX384 thermal cycler (Bio-Rad, Hercules, CA) under the following cycling procedure: 95 °C for 2 min followed by 35 cycles of 95 °C for 10 s, 60 °C for 10 s, and 72 °C for 10 s. Standard curves were generated via 10-fold serial dilutions of genomic *Escherichia coli* DNA (ATTC, MG1655).

#### 2.3.3. 16S sequencing and analysis

For 16S rRNA gene amplicon sequencing, DNA from pooled density fractions from samples labeled with H_2_^18^O and the control (5 per sample-label combination for a total of 330) was amplified in triplicate in 10-µL reactions using primers 515f and 806r [43]. Each reaction contained 1 µL of sample and 9 µL of Phusion Hot Start II High Fidelity master mix (Thermo Fisher Scientific, Waltham, MA) containing 1.5 mM MgCl2. The PCR conditions were as follows: 95 °C for 2 min followed by 20 cycles of 95 °C for 30 s, 64.5 °C for 30 s, and 72 °C for 15 s. The triplicate PCR products were then pooled, diluted 10x, and used as a template in a subsequent dual indexing reaction that used the same primers as above, which also included the Illumina flow cell adaptor sequences and 8-nucleotide Golay barcodes (15 cycles identical to the initial amplification conditions). The PCR conditions were identical to those described above but included 15 cycles. Amplicons were purified with AMPure XP magnetic beads (Beckman Coulter, Brea, CA) and quantified with a PicoGreen assay (Invitrogen, Waltham, MA) on a BioTek Synergy HT plate reader (BioTek Technologies, Vinooski, VT). The samples were pooled at equimolar concentrations, purified with AMPure XP beads (Beckman Coulter, Brea, CA), and quantified using the KAPA Sybr Fast qPCR kit (Kapa Biosciences, Willmington, MA). Libraries were sequenced on an Illumina MiSeq (Illumina, San Diego, CA) at Northern Arizona University’s Genetics Core Facility using a 300-cycle v2 reagent kit.

Demultiplexed reads were right trimmed to 140 nucleotides and quality filtered with DADA2 v1.14.1 (parameters maxN = 0, maxEE = c(2, 2), truncQ = 2) [44]. Chimeric sequences were predicted *de novo* and removed with the removeBimeraDenovo() function in DADA2 using the “consensus” method. The resulting amplicon sequence variant (ASV) table was filtered to keep ASVs with lengths 252-254. Taxonomy was assigned with the RDP classifier v2.11 [45] against training set 16.

#### 2.3.4. Taxon-specific growth and mortality rate calculations

Atom fraction excess (AFE) and growth, mortality, and net rates were calculated using quantitative isotope probing (qSIP) in R (v.4.4.0) [46], as previously described [46]. Specifically, we estimated the number of 16S rRNA gene copies per taxon in each pooled density fraction by multiplying the sequencing-derived taxon-specific relative abundances by the total number of 16S gene copies as measured by qPCR. To maximize the available information on the densities of taxa in the natural abundance isotope treatment, we combined taxon-specific density estimates across the 3 h, 24 h, 48 h, 72 h, and 168 h time points for a total of 15 replicates. We limited our analyses to only those taxa that occurred in at least 3 (of 15) replicates of the natural abundance treatment and in all 3 replicates of the ^18^O treatment to reduce the likelihood of spurious density shifts for rare taxa. Density shifts and atom fraction excess estimates, along with 95% confidence intervals, were computed by bootstrapping (1000 iterations; [47]).

We estimated absolute population growth and mortality rates (# [g-soil]^-1^ d^-1^) as the changes in 16S gene copies per gram soil per time, over each of the time intervals of the incubation: 0-3 h, 0-24 h, 0-48 h, 0-72 h, and 0-168 h. We assumed linear population growth over the course of the incubations, and we estimated taxon-specific net absolute population growth rates and taxon-specific absolute mortality rates for the five time intervals using the equations previously described [47]. The fraction of oxygen in DNA derived from water (versus other organic sources) was assumed to be 0.60 [48]. Uncertainty of growth and mortality rates (95% confidence interval) was estimated using a bootstrapping procedure with 1000 iterations [48]. As highlighted in previous work [48], calculated absolute abundances should be interpreted as approximations, given the various challenges and biases inherent to the entire work flow. These include incomplete extraction of DNA from soil microorganisms [49], differences in 16S rRNA gene copy numbers among taxa, and biases introduced during amplification and sequencing [50, 51]. Consequently, we emphasize that our calculations are best viewed as rough approximations for assessing taxon-specific growth and mortality rates.

#### 2.3.5. Bulk 16S rRNA gene amplicon analysis

We used the sequence data per fraction to approximate the bulk microbial communities at each timepoint. To this end, we first determined how much of the total DNA was represented in each fraction by dividing the 16S RNA gene copy number from each fraction by the total across all fractions. Next, we multiplied the total counts across all fractions by the proportion of DNA represented in each fraction to obtain an adjusted new total count. Finally, the relative abundance of each ASV was multiplied by the total count for each fraction to obtain an adjusted new count for each ASV. This allowed us to sum the counts of each ASV across all fractions to obtain a single value for each sample to obtain a count table. Downstream taxonomic analysis and bar chart creation were performed via the phyloseq R package (v. 1.48.0) [52]. Taxa with relative abundances less than 0.01% were filtered out.

#### 2.3.6. Proxy or Metric for Community growth efficiency (CGE)

To provide a quantitative assessment of carbon allocated to new growth relative to CO_2_ production, the community growth efficiency (CGE) for each timepoint (t) was calculated from net growth (*Gn*) and respiration (R; CO_2_ efflux) as follows, where CGE_max_ was the highest CGE across all samples:

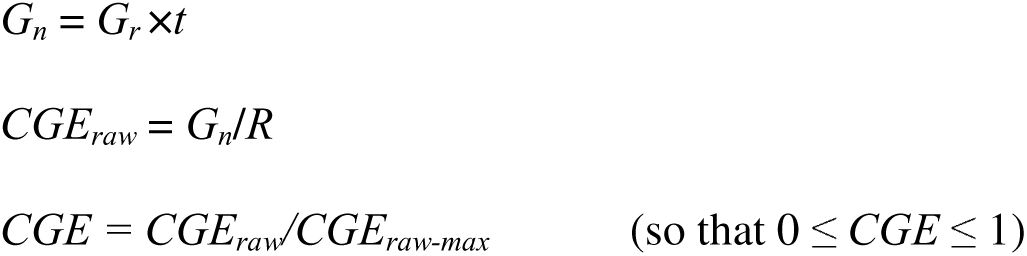

#### 2.3.4. Metagenomic sequencing and analysis

Metagenomic analysis was performed on 9 samples from the 50% MAP treatment (3 plots at 0 h, 48 h, and 168 h; no samples from 24 h or 72 h were sequenced) and 15 samples from the 100% MAP treatment (3 plots at 0 h, 24 h, 48 h, 72 h, and 168 h). For each sample, unfractionated DNA and DNA from 5 density fractions per sample (total sample-fractions = 234) were sequenced following a 2L×L151 indexed run recipe on an Illumina NovaSeq S4 platform (Illumina, San Diego, CA) by Novogene Co. (novogene.com). For metagenomics analysis, Illumina adapters and phiX sequences were removed from raw reads using BBmap (v. 38.79), which is included in the BBtools package [53], and the remaining reads were quality trimmed using Sickle (v. 1.33) [54] for paired-end reads. This process yielded 1.55×10^8^ ± 1.5×10^7^ (mean ± standard deviation) reads per sample. For unfractionated samples, reads from triplicate samples (plots) from each time point were coassembled, and for fractionated samples, reads from all fractions within a sample were coassembled using MEGAHIT (v. 1.2.9) [55] with presets of “large-meta” and a minimum contig length of 1000 bp. DRAM (v. 1.5.0) [56] was used for gene prediction and annotation using KEGG orthology (KO) using Hidden Markov Models (HMM) for all KO entries within the KOfam database [57], and with the carbohydrate active enzyme (CAZy) entries within dbcan [58]. A reference gene catalog was created by clustering the amino acid sequences of all the predicted genes across all the unfractionated and fractionated coassemblies at the 95% confidence level using Mmseq2 (v. 13.45111) [59] to create a dereplicated list of genes to map metatranscriptome reads.

### 2.4. RNA extraction and metatranscriptome sequencing and analysis

Metatranscriptomics was performed on all 48 samples (2 legacy precipitation treatments × 3 plots × 6 timepoints), and a full description of RNA extraction and sequencing is provided in [37, 60]. Briefly, RNA was extracted from soils via the RNeasy PowerSoil Total RNA Kit (Qiagen, Hilden, Germany) according to the manufacturer’s instructions. Extracted RNA was treated with RNase-free DNase (Qiagen, Hilden, Germany) and stored at −80 °C. The RNA concentration was measured with a Qubit fluorometer (Invitrogen, Waltham, MA), and the quality was assessed with a Nanodrop One Spectrophotometer (Thermo Fisher Scientific, Waltham, MA). The samples were sent to the Joint Genome Institute (JGI; Berkeley, California) for rRNA depletion, library preparation, and sequencing. Ribosomal (r) RNA was removed from 100 ng of total RNA via Qiagen FastSelect 5S/16S/23S for bacterial rRNA depletion (and additional FastSelect plant and/or yeast rRNA depletion) (Qiagen, Hilden, Germany) with RNA blocking oligo technology. One heavily degraded sample was discarded (P04_x50_t0; t0 sample from Plot 4 of 50% legacy moisture treatment), and the remaining 47 samples were reverse transcribed to create first-strand complementary (c) DNA using the Illumina TruSeq Stranded mRNA Library prep kit (Illumina, San Diego, CA), followed by second-strand cDNA synthesis, which incorporated dUTP to quench the second strand during amplification. The double-stranded cDNA fragments were then A-tailed and ligated to JGI dual-indexed Y-adapters, followed by enrichment of the library by 10 cycles of PCR. The prepared libraries were quantified via a KAPA Biosystems next-generation sequencing library qPCR kit (Kapa Biosciences, Willmington, MA) and run on a Roche LightCycler 480 real-time PCR instrument. Sequencing of the flow cell was performed on the Illumina NovaSeq sequencer (Illumina, San Diego, CA) using NovaSeq XP V1.5 reagent kits and an S4 flow cell, following a 2L×L151 indexed run recipe.

The raw reads were downloaded from the JGI genome portal and quality control filtered via bbduk (from the BBtools [v. 38.79] package [53]) and Sickle (v. 1.33) [54]. rRNA was then filtered from the paired-end reads via SortMeRNA (v. 4.3.6) [61], followed by repair.sh (BBtools [v. 39.06] package) to correct the read orientation. After all the filtering steps, 31.6×10^6^ ± 1.5×10^6^ reads per sample remained. Reads were mapped via BBmap (BBtools [v. 39.06]) to the metagenome reference gene catalog, and counts and transcripts per million (tpm) were calculated via coverM (v. 0.70) [62]. The mapping rate was 23.2 ± 8.4% (except for P08_x100_t3 which had no reads mapped and was removed from downstream analyses). The taxonomies of the reads were determined via Kraken 2 (v. 2.1.3) [63]. Normalization of counts was performed to obtain variance stabilized transformations (VSTs) using DESeq2 (v. 1.44.0) [64].

### 2.5 Fourier transform ion cyclotron resonance mass spectrometry (FTICR-MS)

For each sample, 1 g of lyophilized soil was defrosted, added to a clean tube, and extracted with 2 mL of Milli-Q water (>18.2 MΩ·cm resistivity). The samples were shaken at 1000 rpm at room temperature for 2 hours on a vortex shaker and then centrifuged at 6000 rpm for 5 minutes, after which the supernatant was removed. This process was repeated with another 2 mL of Milli-Q water, and the supernatants were combined. Solid phase extraction (SPE) was performed on water extracts using Bond Elut Priority Pollutant (PPL) cartridges (Agilent Technologies, Santa Clara, CA) to remove salts and impurities that could interfere with mass spectrometry (MS) analysis [65]. Water extracts were diluted to 5 mL with Milli-Q water and adjusted to pH 2 with ∼2 µL of 85% H_3_PO_4_ prior to addition to methanol-activated PPL cartridges. The organic matter bound to the cartridges was rinsed with 50 mL of 10 mM HCl, dried with nitrogen and eluted in 1.5 mL of methanol. Following the water extraction, a modified metabolite, protein, lipid extraction (MPLEx) [66, 67] was performed to yield polar and nonpolar liquid fractions. Ice-cold chloroform and methanol (2:1, total ×6 mL) were added to the soil, which was vortexed after each addition. Water was added (0.25 mL), and the samples were shaken for 1 hr at 1000 RPM. Another 1.25 mL of water was added at a final solvent ratio of 8:4:3 (chloroform:methanol:water), and the samples were gently shaken before being incubated at 4 °C overnight. The samples were then centrifuged at 6000 RPM for 5 minutes, yielding two layers. The layers were removed separately and frozen at −80 °C until individual analysis.

Fourier transform ion cyclotron resonance (FTICR)-MS data were acquired using a 7 T Bruker ScimaX FTICR-MS (Bruker, Billerica, MA) operated with quadripolar (2×) detection and located in the Environmental Molecular Science Laboratory at Pacific Northwest National Laboratory (Richland, WA, USA). External calibration was performed with sodium trifluoracetic acid followed by shimming of the magnet to minimize 1× harmonic resonance peaks and tuning to optimize the spectra for peak intensity, shape and resolution over the m/z range of 200–1000. The extracts were directly infused into the electrospray source in negative ion mode at a voltage of +4 kV, a temperature of 200 °C, and a dry gas flow of 4 L/min in randomized order via a custom automated cart. The ion accumulation time was set to 10 ms, and 2.1 s transients were added over 300 acquisitions in an 8 MW time domain for an estimated average resolution of ∼680 K at m/z 400. Internal recalibration of each spectrum, post-acquisition, was performed with calibration lists of standard organic matter (OM) components. Peak picking was performed during data analysis using a signal-to-noise ratio (S/N) threshold of 7, a relative intensity threshold of 0.01, and an absolute intensity threshold of 100,000.

The peak lists were exported, and Formularity software [68] was used to align the spectra within a 0.5 ppm threshold and assign formulas of N ≤ 2, S = 0 and P = 0, and <0.5 ppm error for high confidence assignments (average error <0.2 ppm). The peak lists were then corrected by subtracting the peaks detected in the extraction blanks. Elemental ratios were calculated from the molecular formulas assigned for each peak and averaged for each sample.

Analysis of the FTICR data was performed using MetaboDirect (v.0.2.7) [69] to 1) create profiles of biochemical compound classes for each of the thousands of peaks in each sample’s electrospray ionization (ESI) FTICR-MS spectrum using van Krevelen diagrams and 2) calculate thermodynamic parameters (nominal oxidation state of carbon [NOSC] and Gibb’s free energy [GFE]). Each peak was assigned a biochemical compound class on the basis of its predicted formula and plotting of C, H, and O on a van Krevelen diagram as follows: s (0<C:C≤0.3 and 1.5≤H:C≤2.5), unsaturated hydrocarbons (0≤O:C≤0.125 and 0.8≤H:C≤2.5), proteins (0.3<O:C<0.55 and 1.5≤H:C≤2.3), amino sugars (0.55<O:C≤0.7 and 1.5≤H:C≤2.2), lignin (0.125<O:C≤0.65 and 0.8≤H:C<1.5), tannins (0.6<O:C≤1.1 and 0.8≤H:C<1.5) and condensed hydrocarbons (aromatics; O≤200 O:C≤0.95 and 0.2≤H:C<0.8) [70]. This method is semiquantitative, and each compound class is then reported as relative abundance values on the basis of counts (presence/absence). For the methanol extraction, two outliers were removed: P10_x50_t72 and P13_x50_t71.

### 2.6 Statistical analysis

All the statistical tests were performed in R v. 4.4 [46].

#### 2.6.1 Linear mixed effects models

Significant differences in the NOSC and GFE of unique compounds as well as the growth/mortality rate, net growth, 16S rRNA gene abundance (biomass), bulk 16SrRNA phyla, and CO_2_ efflux between the legacy precipitation treatments over time post rewet were assessed by fitting a linear mixed-effects (LME) model. LME models were created using the lmer() function in the lmerTest package [71] with the following formula:

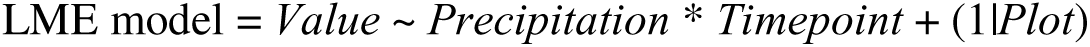

where *Value* is the measurement, *Precipitation* (50% vs 100%) and *Timepoint* (0 h, 3 h, 24 h, 48 h, 72 h, and 168 h) are the fixed effects, and *Plot* (location sample was collected from) is the random intercept to account for repeated measures. To determine the significance of each fixed effects term in the LME model, an analysis of variance (ANOVA) was performed using the anova() function included in base R. To test for significant differences at each timepoint, we computed estimated marginal means (least-squares means) and Tukey’s honestly significant difference (HSD)-adjusted pairwise comparisons with the emmeans() and pairs() functions in the emmeans package (v. 1.11.1) [72].

#### 2.6.2 G test for unique compounds

To test for the uniqueness of the FTICR compounds between the legacy precipitation treatments, we used a G test of independence. First, we extracted all formulas that were detected in only the 50% or 100% legacy precipitation treatments, where a compound was considered detected if it was present in at least two of the three plots and determined the counts for each compound class (amino sugar-, carbohydrate-, condensed hydrocarbon-, lignin-, protein-, tannin-, and unsaturated hydrocarbon-like compound). We tested for a shift in unique compound class composition between treatments with a log-likelihood ratio (G-) test using the GTest() function in the DescTools package (v. 0.99.60) [73]. We then performed a G test on each compound class individually (with all other compounds in the “other” category) to determine a G value and associated p value per class. False discovery rate (FDR)-corrected p values are reported.

#### 2.7.3 Nonmetric multidimensional scaling

Nonmetric multidimensional scaling (NMDS) was performed to identify relationships between samples and identify major drivers for each meta-omics dataset (FTICR compounds [WEOC and MEOC] and transcript abundances [metatranscriptomics] at the KO functional level). Analysis was performed on BrayLCurtis dissimilarity matrices for transcript abundances and Jaccard distance matrices for FTICR using the metaMDS function in the vegan package (v. 2.7-1) [74]. PERMANOVA was used to determine significant differences between treatment groups (i.e., timepoint relative to wet-up and legacy precipitation) using the adonis2 function in the vegan package.

#### 2.7.4 Partial least square regression

Partial least squares regression (PLSR) was used to identify genes whose related metabolic pathways were most strongly associated with CGE. As predictor variables, the VST counts of all KOs and the CAZy enzyme class were used across samples from all timepoints, except for 0 h. The response variable was the log-transformed CGE metric. We selected the genes with variable importance in projection (VIP) scores in the top 10% as those that contributed most to CGE. The direction of correlation was determined by whether the coefficient was positive or negative. KO genes were mapped onto KEGG metabolic pathways using the online tool Interactive Pathway Explorer v3 (https://pathways.embl.de/).

#### 2.7.5 Correlations

Pearson correlations were calculated between the log-transformed CGE metric and the transcript abundances of CGE-associated genes (VST counts) within specific biosynthesis and degradation pathways for key biomolecules and energy-related pathways from curated lists (**Table S1**). The specific pathways within each of these categories are as follows: 1) the biosynthesis list included amino acid, lipid, nucleotide, peptidoglycan, teichoic acid, and extracellular polymeric substance [EPS] biosynthesis pathways; 2) the degradation lists included amino acid, lipid, nucleotide, aromatic, and C-ring (subset of aromatic) degradation pathways; and 3) the energy-related lists included central C metabolism, pyruvate oxidation (subset of central metabolism), electron transport chain, adenosine triphosphate production (ATP-ase), fermentation, and the reactions between phosphoenolpyruvate-oxalacetate-malate-pyruvate (POMP). Correlations were performed between the sum of transcript abundances (VST counts) of all KOs within each pathway and the log-transformed CGE metric. Linear regressions were performed to visualize relationship between VST counts and CGE metric on plots and Pearson correlations to determine degree and significance of correlation with r and p-values (FDR-corrected).

## 3. Results

### 3.1 Growth, microbial turnover and community growth efficiency (CGE)

Large differences in taxon-specific growth rates were evident between legacy precipitation regimes (**Fig. 1a-d**). At 3 h post rewatering, the normal legacy precipitation (100% MAP) treatment was dominated by the growth of *Proteobacteria*, *Firmicutes*, and *Actinobacteria* and the mortality of *Proteobacteria* and *Actinobacteria*, in decreasing order of rates (**Fig. 1a, b**). In contrast, growth under reduced legacy precipitation (50% MAP) was dominated by *Firmicutes*, *Proteobacteria*, and *Actinobacteria* in decreasing order of rates, and mortality was dominated by *Actinobacteria* (**Fig. 1c, d**). All taxa at the phylum level presented the highest growth rates at 3 h across both legacy precipitation regimes, except for *Proteobacteria* at reduced precipitation, which presented the highest growth rate at 24 h post rewatering. This finding reflects the differences in the bulk communities representative of all active and non-active microbes, where reduced legacy precipitation had a greater overall relative abundance of *Actinobacteria* across all timepoints (p=3e-9; df=1; LME) and specifically at 0h before rewetting (p=4e-4, Tukey HSD; **Fig. 1e**). Normal legacy precipitation had a greater overall relative abundance of *Proteobacteria* across all timepoints (p=0.01, df = 1, LME; **Fig. 1e**) and specifically at 0 h (p=0.002; Tukey HSD). Despite a greater proportion of *Firmicutes* growing after rewetting in the reduced legacy precipitation treatment, the relative abundance of *Firmicutes* at the bulk level across all timepoints, including before rewetting at 0 h, did not differ between the legacy precipitation treatments (p>0.05, LME and Tukey HSD; **Fig. 1e**). These shifts in microbial communities at the bulk level were also evident in beta diversity, where legacy precipitation treatment was a significant driver of community composition (p=0.001, df = 1, PERMANOVA).

**Figure 1.**
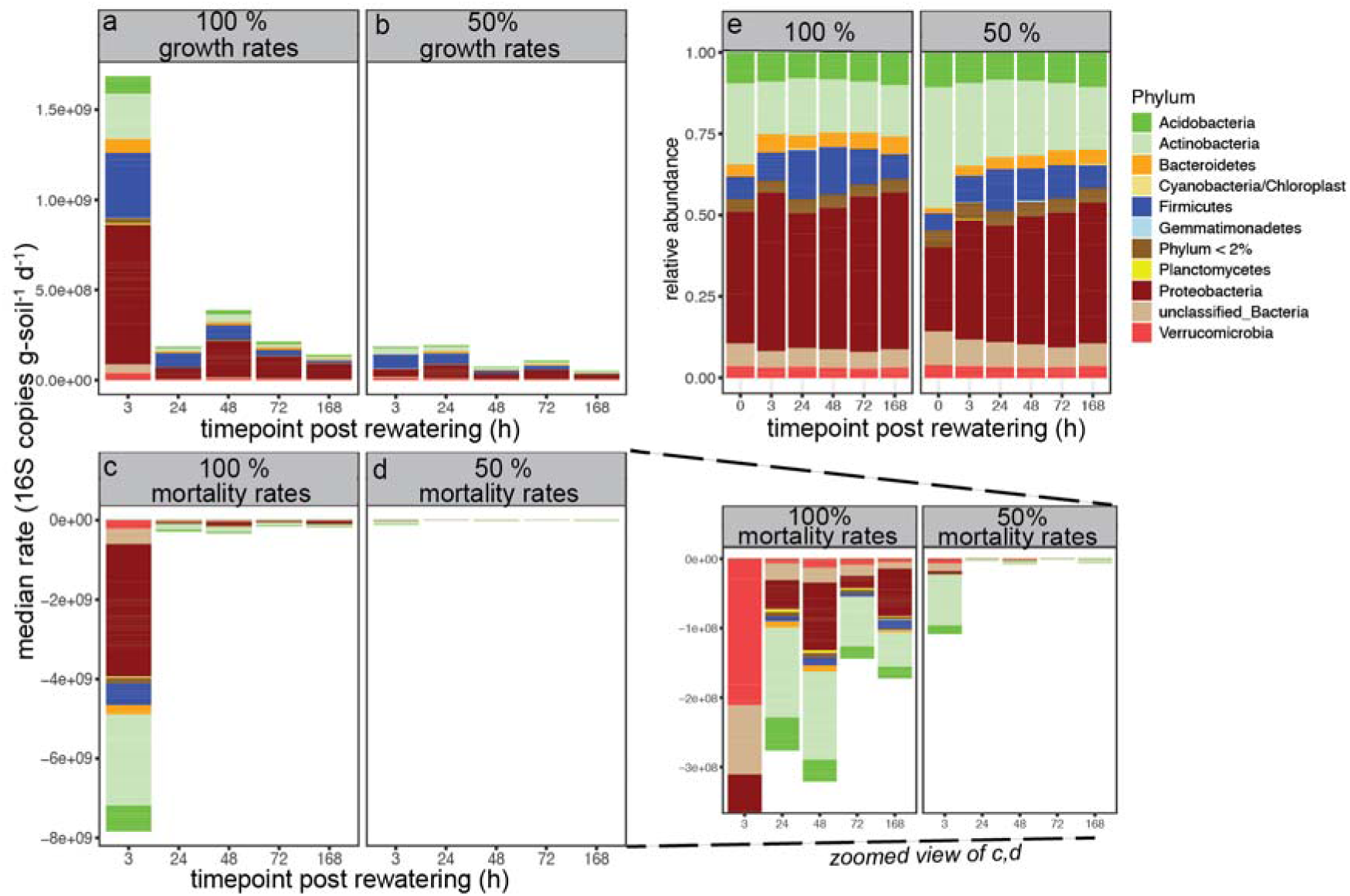
Taxon-specific growth and mortality rates and bulk community composition. a) Growth rates for normal (100% MAP) and b) reduced (50% MAP) legacy precipitation. c) Mortality rates for normal and d) reduced legacy precipitation. e) Bulk changes in community composition as determined with 16S rRNA gene amplicon sequencing. Colors across panels indicate phylum-level classifications of taxa.

These overall patterns in growth dynamics were reflected at the community level, where growth and mortality rates across timepoints were significantly higher for the normal than for the reduced legacy precipitation treatments (p=4e-14 and p=8e-15, respectively, df=4, LME; **Fig. 2b, c**). The highest growth and mortality rates under normal legacy precipitation occurred at 3 h and were 2.1×10^9^ and 9.5×10^9^ 16S copies g-soil^-1^ d^-1^, respectively, and the highest growth and mortality rates under reduced legacy precipitation occurred at 24 h and 3 h and were 2.3×10^8^ and 1.2×10^8^ 16S copies g-soil^-1^ d^-1^, respectively.

**Figure 2.**
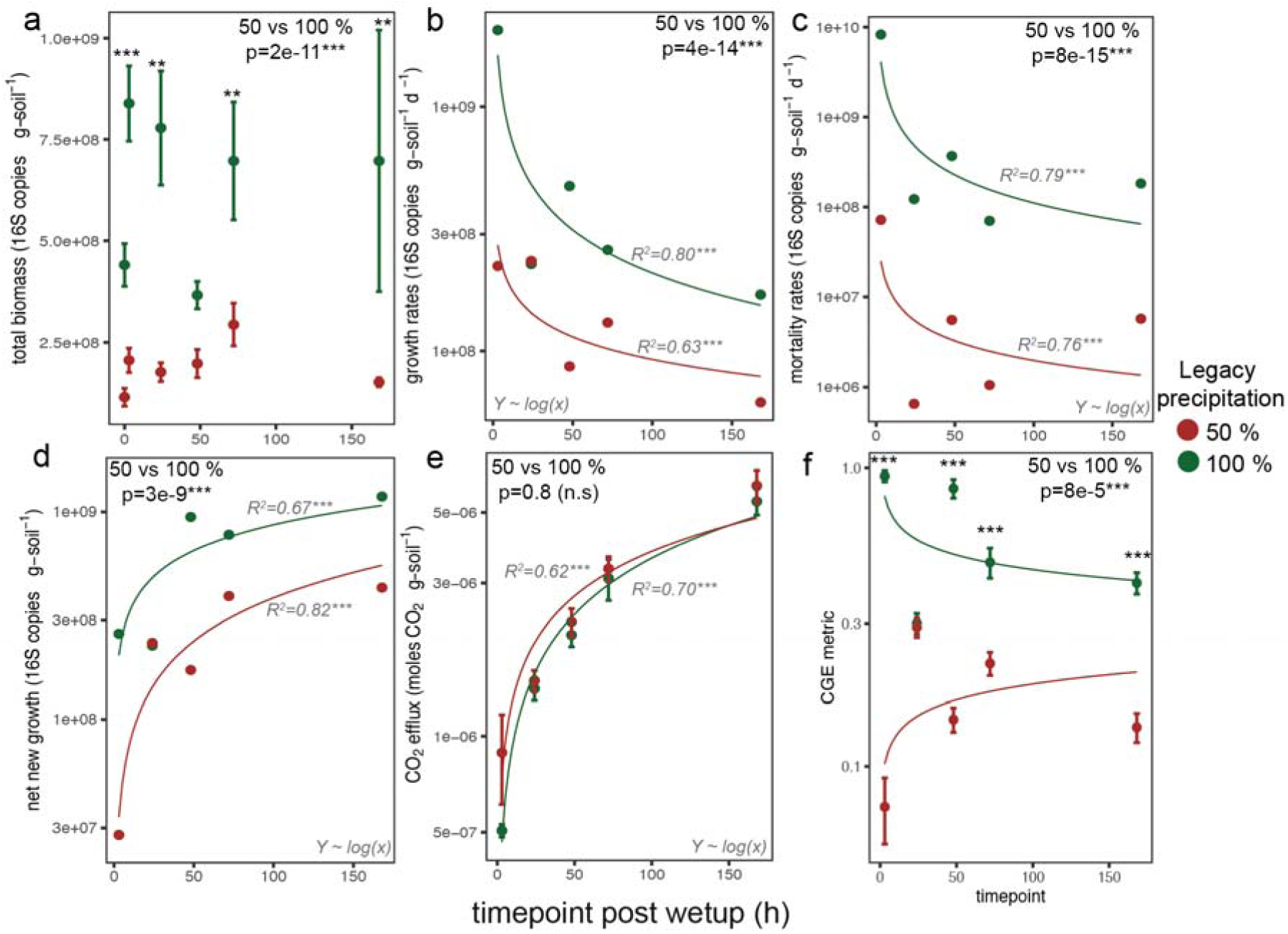
Microbial growth dynamics upon rewatering post summer dry down for normal (100% MAP) and reduced (50% MAP) legacy precipitation. a) Total biomass calculated from qPCR of 16S rRNA gene and representing both active and nonactive bacteria and archaea. b) Growth rates and c) morality rates. d) Cumulative biomass and e) cumulative CO_2_ efflux at each time point post rewatering. f) Community growth efficiency (CGE) metric measured as ratio of new biomass to cumulative CO_2_ efflux, normalized to a value between 0 and 1. Differences between legacy precipitation treatments were assessed with Linear Mixed-Effects models (panels a-f) followed by Tukey HSD to compare treatment at each time point (panel a, f). Linear regressions for each legacy moisture treatment were performed to capture linear trend over time (panels b-e) with best fit shown and R^2^. For panel f, linear trend shown but R^2^ were not significant for 50% legacy precipitation. *, p < 0.05; **, p < 0.01; ***, p < 0.001

The total biomass, measured as the total 16S rRNA gene abundance as determined by qPCR and representing both active and inactive bacteria and archaea, was overall greater in the normal than in the reduced legacy precipitation treatments (p=2e-11; df=4, LME; **Fig. 2a**). While the cumulative new biomass, which was calculated from growth rates, was greater under normal compared to reduced legacy precipitation (p=3e-9, df=4, LME; **Fig. 2d**), the CO_2_ efflux was similar in both treatments (p=0.4, df=4, LME; **Fig. 2e**). This resulted in a divergence in CGE, which was greater in the normal compared to the reduced legacy precipitation treatments (p=8e-5, df=4, LME; **Fig. 2f**), indicating a decline in CGE metric at reduced legacy precipitation associated with an increase in soil CO_2_ loss per unit of new growth.

### 3.2 Dynamics of meta-omics driven by rewatering and legacy precipitation

After observing the drastic impact of legacy precipitation on microbial communities, community growth rates, and CGE, we sought to determine the impact of legacy precipitation on changes in SOC, measured as water-extractable organic carbon (WEOC) and methanol-extractable organic carbon (MEOC) from FTICR-MS, and gene transcript abundances (metatranscriptomics). For these datasets, timepoint was the primary driver of changing SOC composition and gene expression (WEOC [p<0.001], MEOC [p=0.01], gene expression [p<0.001; df=5; PERMANOVA), whereas legacy precipitation was a secondary driver for only WEOC (p=0.006) and MEOC (p=0.01, df=1, PERMANOVA) (**Fig. S1**).

#### 3.2.1 Water-extractable organic carbon (WEOC)

The composition and thermodynamics of soil WEOC, which is indicative of dissolved organic matter readily accessible to microbes, differed between timepoints and legacy precipitation. In total, 119,705 and 122,285 compounds were detected in the reduced and normal legacy precipitation treatments, respectively, and of these, 8,660 and 8,307 from the reduced and normal legacy precipitation treatment, respectively, were assigned formulas and classified into compound classes on the basis of their O:C and H:C ratios. Class-assigned compounds, averaged across legacy precipitation treatments, were characterized as follows: lignin-like compounds dominated with an average proportion of 47.9%, followed by condensed hydrocarbon-like compounds (32.4%), tannin-like compounds (14.5%), protein-like compounds (2.0%), carbohydrate-like compounds (1.0%), lipid-like compounds (1.0%), amino sugar-like compounds (0.8%), and unsaturated hydrocarbon-like compounds (0.2%) (**Fig. S2c)**.

A comparison of the reduced and normal legacy precipitation treatments revealed that there were 643 compounds were only present in the reduced legacy precipitation treatment (“unique” to reduced precipitation), and 280 compounds were only present in the normal legacy precipitation treatment (“unique” to normal precipitation). Within these unique compounds, there was a significant shift in class-level categorization (p<2eL¹L, G=229, df=8, G test). In the reduced legacy precipitation treatment, tannin- and lignin-like compounds, which are characteristic of decomposing plant detritus, were overrepresented (p=4e-10, G=41, and p=6e-4, G=12, respectively; G test), as were condensed hydrocarbons (p=4e-6, G=22.4, G test). In the normal legacy precipitation treatment, protein-like (p=9e-11, G=45, G-test), lipid-like (p=1e-7, G=30, G-test), carbohydrate-like (p=1e-3, G=11, G-test), and amino sugar-like (p=0, G=71, G-test) compounds, characteristic of necromass, were overrepresented, as were unsaturated hydrocarbon-like compounds (p=3e-5, G=18, G-test) (**Fig 3a, b**; see **Fig S3d** for the van Krevelen diagram with all the compounds). This shift in the class composition of compounds unique to normal and reduced legacy precipitation was associated with decreased mean Gibb’s free energy (GFE) and increased nominal oxidation state of carbon (NOSC) (p<2e-16, df=1, LME; **Fig. 3c, d**).

**Figure 3.**
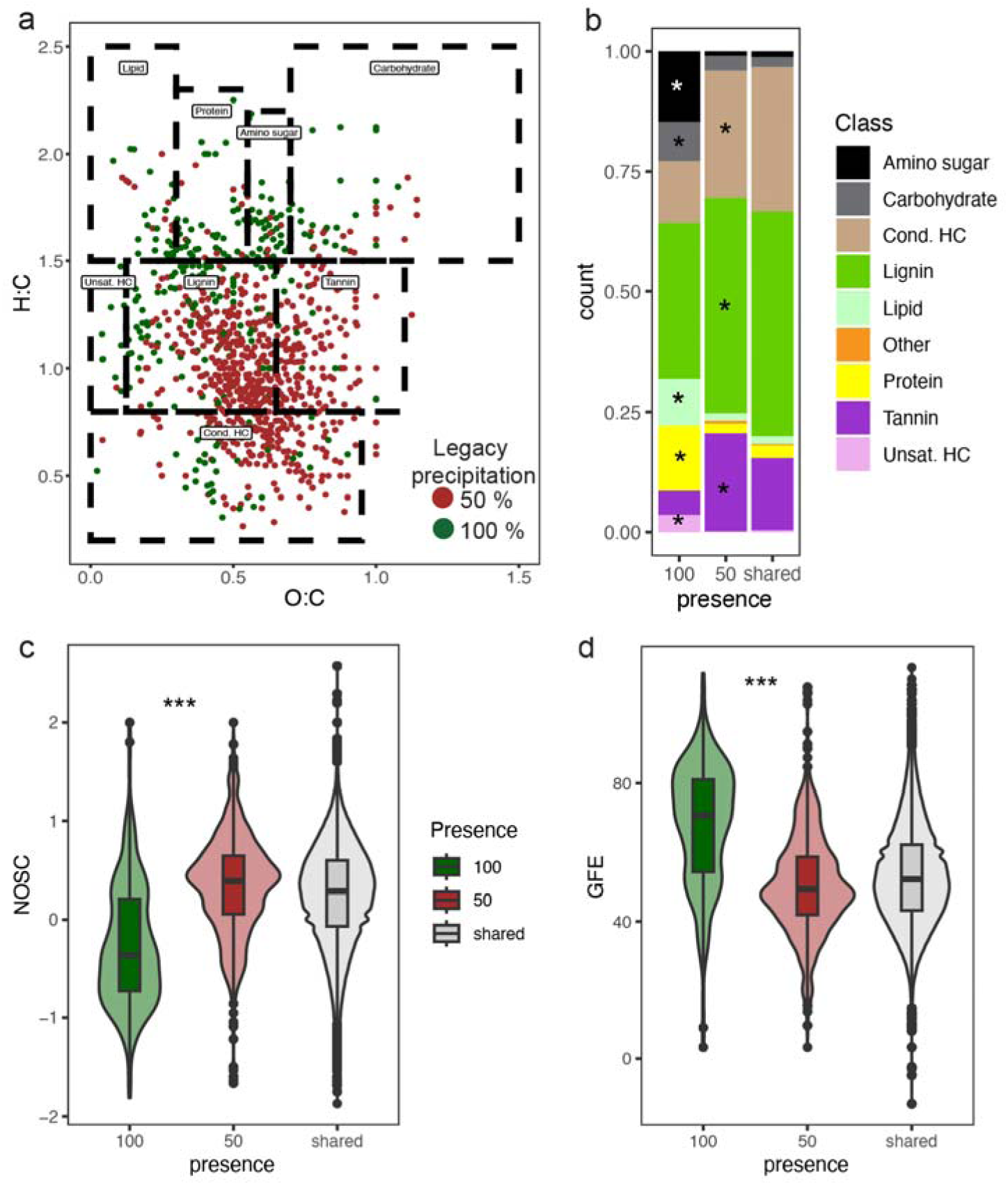
Soil water extractable organic carbon (WEOC) differs between normal (100% MAP) and reduced (50% MAP) legacy precipitation. A) Van Krevelen Diagram showing unique compounds (only compounds in either 50 or 100% MAP). Ratios of O:C vs H:C allow for predictions of compound classes (showed in doted square boxes). b) Relative abundances of compound classes of unique compounds found in 100%-only and 50%-only as well as for compounds found in both 50 and 100% legacy precipitation treatments. Compounds with a star are significantly overrepresented in respective treatment (G-test). (c-d) Violin plots and embedded boxplots (represent Q1-Q3, centerlines indicate the median and whiskers extend to the minimum and maximum values, excluding outliers [black points]) showing mean and distribution of c) nominal oxidation state of carbon (NOSC) and d) Gibbs free energy (GFE) for unique compounds within each precipitation treatment. For c-d: ***, p<0.001 (unique compounds in normal vs reduced precipitation, LME, df = 1)

At the class level, there were trends across time after rewetting (**Fig. S3a**). Carbohydrate-like and lipid-like compounds decreased in proportion in both the reduced and normal legacy precipitation treatments over time between 0 h and 168 h (p<0.05 and p<0.001, respectively; df=5; Tukey HSD). Amino sugar-like compounds decreased in proportion in only the normal legacy precipitation treatment between 0 h and 168 h (p<0.05, df=5, Tukey HSD). Condensed hydrocarbon-like compounds decreased in proportion in both the reduced and normal legacy precipitation treatments between 0 h and 168 h (p<0.05, df=5, Tukey HSD) and unsaturated hydrocarbons decreased in only the normal legacy precipitation treatment between 0 h and 168 h (p<0.001, df=5, Tukey HSD). Lignins and tannins did not change significantly over time.

#### 3.3.2 Methanol extractable organic carbon (MEOC)

The composition and thermodynamics of soil MEOC, which is indicative of more polar organic matter that is not as readily accessible to microbes as WEOC is, also differed across timepoints post rewetting and legacy precipitation. In total, 51,829 and 55,963 compounds were detected in the reduced and normal legacy precipitation treatments, respectively, and of these, 5,358 and 5340 from the reduced and normal legacy precipitation treatment, respectively, were assigned formulas and classified into compound classes. Class-assigned compounds across legacy precipitation treatments were characterized as follows: lignin-like compounds dominated with an average proportion of 38.9%, followed by condensed hydrocarbon-like compounds (17.1%), protein-like compounds (15.2%), carbohydrate-like compounds (11.9%), amino sugar-like compounds (8.7%), lipid-like compounds (5.7%), tannin-like compounds (2.1%), and unsaturated hydrocarbon-like compounds (0.3%) (**Fig. S3e**). Compared with the WEOC, there were more protein-like, carbohydrate-like, amino sugar-like, and lipid-like MEOCs and fewer tannin-like and condensed hydrocarbon-like MEOC compounds. Compounds such as amino sugars and proteins are often considered polar; however, they may contain hydrophobic regions that can sorb to soil minerals, making them less bioavailable [75, 76].

There were 725 compounds only present in the reduced legacy precipitation treatment (“unique” to reduced precipitation), and 293 compounds were only present in the normal legacy precipitation treatment (“unique” to normal precipitation). Among these unique compounds, there was a smaller shift in class-level categorization between legacy precipitation treatments than in the WEOC, but this shift was still significant (p<2eL¹L, G=87, df=8, G test) with carbohydrate-like compounds being enriched in the normal legacy precipitation treatment (p=5e-4, G=16) (**Fig. S2a, b**; see **Fig. S3f** for the van Krevelen diagram with all the compounds). Similar to WEOC, this shift in the class composition of unique compounds from normal to reduced legacy precipitation was associated with a lower mean GFE and higher NOSC (p=3e-7, df=1, LME**; Fig. S2c, d**).

At the class level, there were fewer trends across time post laboratory rewet than WEOC, with the largest shift occurring between pre- (0 h) and post rewet (all other timepoints), particularly with a decrease in amino sugars and carbohydrates at normal precipitation between 0 and 3 h (p=0.02, df=18, Tukey HSD) (**Fig. S3b**). Furthermore, at 0 h, there was a greater abundance of amino sugars and carbohydrates in the normal-precipitation treatment than in the reduced-precipitation treatment (p=0.02 and 0.05, respectively; df=18; Tukey HSD) and a greater abundance of condensed hydrocarbons in the reduced-precipitation than in the normal precipitation (p=0.04, df=18, Tukey HSD; **Fig S3b**). Compared with those at 0 h, the abundance of condensed hydrocarbons tended to increase with time at 48, 72, and 168 h post rewetting (p=0.02, p=0.03, and p=0.04, respectively; df=18; Tukey HSD).

### 3.3. Gene expression and community growth efficiency metric

Next, we identified genes and their metabolic pathways that were most associated with CGE metric using PLSR. We screened all the KO-annotated genes and identified 644 that were associated with CGE (top 10% of the VIP scores ranging from 1.6–3.5) (**Table S4**). Among these genes, 389 were positively associated with CGE and 254 were negatively associated with CGE. Transcripts for genes associated positively with CGE metric were involved in amino acid metabolism (biosynthesis of threonine, methionine, lysine, and tryptophan, serine), nucleotide metabolism (biosynthesis and degradation of purines and pyrimidines), xenobiotics metabolism (degradation of benzoate and catechol) pathways, and energy (TCA cycle, oxidative phosphorylation) (**Fig. 4a**), whereas transcription associated negatively with CGE metric were associated with amino acid metabolism (biosynthesis of lysine, serine, and proline; degradation of lysine), lipid metabolism (biosynthesis of fatty acid, glycerolipids, and glycerophospholipids), carbohydrate metabolism (breakdown of trehalose, glycogen, and cellulose and pentose and glucoronate metabolism), nucleotide metabolism (biosynthesis and degradation of purines and pyrimidines), and energy (glycolysis, oxidative phosphorylation, and pentose phosphate pathway) metabolism pathways (**Fig. 4b**).

**Figure 4.**
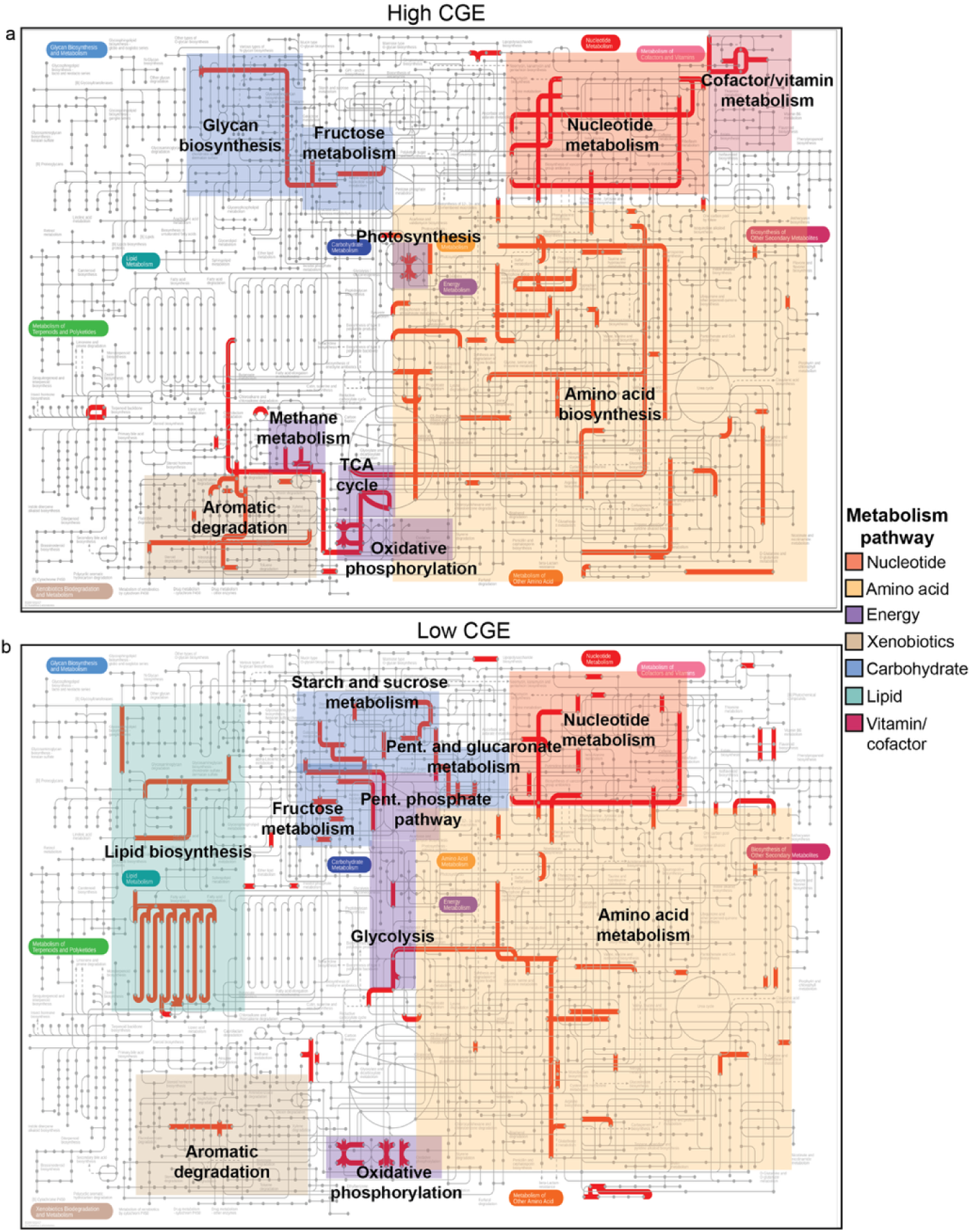
KEGG global metabolic pathways of all genes associated a) positively with community growth efficiency (CGE) and b) negatively associated with CGE. Pathways are highlighted with color based on category and labeled with description. Categories are colored as: Red- nucleotide, tan- xenobiotic, purple- energy, blue- carbohydrate, green- lipid, orange-amino acid, and magenta- vitamin and cofactor metabolism. Pent., pentose; TCA, tricarboxylic acid

Next, we focused on whether the expression of CGE-associated genes within specific biosynthesis, degradation, and energy pathways from our curated lists was correlated with the CGE metric (**Table 1; Table S2**). Within our curated lists we found 74 genes that were associated positively with CGE and of these, 26% were involved in biosynthesis, 64% were involved in degradation, and 11% involved in energy metabolism. We found 53 genes that were associated negatively with CGE and of these, 26% were involved in biosynthesis, 49% were involved in degradation, and 25% were involved in energy metabolism.

**Table 1.** Summary of CGE-associated genes that belong to each pathway and sub-pathway within the curated lists of biosynthesis, degradation, and energy genes.

| Category | Pathway | Sub-pathway (total number of genes) | Genes associated with CGE |  | Sum of genes within each pathway |  |
| --- | --- | --- | --- | --- | --- | --- |
|  |  |  | Negative | Positive | Negative | Positive |
| Biosynthesis | Amino acid | Alanine (8) | 0 | 2 | 5 | 11 |
|  |  | Chorismate (15) | 0 | 1 |  |  |
|  |  | Cysteine (15) | 1 | 1 |  |  |
|  |  | Lysine/methionine (5) | 0 | 1 |  |  |
|  |  | Serine (18) | 2 | 2 |  |  |
|  |  | Tryptophan (14) | 0 | 1 |  |  |
|  |  | Shared/ Other (154) | 2 | 3 |  |  |
|  | Lipid | Fatty acid (38) | 1 | 0 |  |  |
|  |  | Glycerolipid (42) | 1 | 0 | 3 | 0 |
|  |  | Glycerophospholipid (42) | 1 | 0 |  |  |
|  | Nucleotide | Purine (32) | 1 | 0 |  |  |
|  |  | Pyrimidine (15) | 2 | 2 | 4 | 3 |
|  |  | Ribonucleotide (7) | 1 | 1 |  |  |
|  | Peptidoglycan | Aminosugar (22) | 0 | 1 |  |  |
|  |  | Peptidoglycan unit (22) | 1 | 0 | 1 | 2 |
|  |  | Remodeling (21) | 0 | 1 |  |  |
|  | EPS | Acetan/xanthan (12) | 1 | 0 | 1 | 1 |
|  |  | Alginate (7) | 0 | 1 |  |  |
|  | Teichoic acid | from UND-PP to wall teichoic acid (42) | 0 | 2 | 0 | 2 |
| Degradation | Amino acid | Alanine/aspartate/asparagine/glutamate/glutamine (23) | 2 | 1 |  |  |
|  |  | Arginine/Proline (38) | 3 | 3 |  |  |
|  |  | Cysteine/methionine (45) | 1 | 4 |  |  |
|  |  | Lysine (22) | 1 | 0 | 13 | 12 |
|  |  | Phenylalanine/tyrosine/tryptophan (101) | 2 | 4 |  |  |
|  |  | Serine/glycine/threonine (29) | 2 | 0 |  |  |
|  |  | Shared (52) | 2 | 0 |  |  |
|  | Lipid | Other | 1 | 0 | 1 | 0 |
|  | Nucleotide | Purine | 2 | 1 | 4 | 3 |
|  |  | Pyrimidine | 2 | 2 |  |  |
|  | Xenobiotics | Aminobenzoate (62) | 1 | 2 |  |  |
|  |  | Atrazine (16) | 0 | 1 |  |  |
|  |  | Benzoate (52) | 1 | 8 |  |  |
|  |  | Caprolactam (16) | 0 | 1 |  |  |
|  |  | Chlorocyclohexane and chlorobenzene (16) | 0 | 1 |  |  |
|  |  | Furfural (8) | 1 | 0 | 6 | 22 |
|  |  | Naphthalene (18) | 0 | 2 |  |  |
|  |  | Nitrotoluene (18) | 1 | 1 |  |  |
|  |  | Shared (87) | 2 | 4 |  |  |
|  |  | Steroid (15) | 0 | 1 |  |  |
|  |  | Styrene (17) | 0 | 1 |  |  |
|  | Aromatic ring (AR) | Dihydroxylation of AR (6) | 0 | 1 |  |  |
|  |  | Dihydroxylation and meta-cleavage of AR (25) | 1 | 1 |  |  |
|  |  | Ortho-cleavage of dihydroxylated AR (9) | 1 | 2 | 2 | 10 |
|  |  | Ortho-cleavage of halogenated AR (2) | 0 | 1 |  |  |
|  |  | Other pathway (36) | 0 | 4 |  |  |
|  |  | Ring removal from polycyclic AR (12) | 0 | 1 |  |  |
| Energy | ATPase | ATP-binding (75) | 2 | 3 |  |  |
|  |  | F-type (29) | 1 | 0 | 4 | 3 |
|  |  | P-type (14) | 1 | 0 |  |  |
|  | Central C metabolism | Glycolysis/Gluconeogenesis (46) | 2 | 0 |  |  |
|  |  | PPP (16) | 1 | 0 |  |  |
|  |  | Pyruvate oxidation (11) | 1 | 0 | 4 | 1 |
|  |  | TCA cycle (49) | 0 | 1 |  |  |
|  | ETC | Cytochrome C oxidase (41) | 3 | 0 | 4 | 3 |
|  |  | NADH reductase (80) | 1 | 3 |  |  |
|  | Fermentation | to acetate (10) | 0 | 1 | 0 | 1 |
|  | Pyruvate ox. | Pyruvate => acetyl-CoA (29) | 1 | 0 | 1 | 0 |
| **Total sum** |  |  | **53** | **74** |  |  |

CGE-associated genes were determined by partial least squares regression (PLSR) analysis and selecting genes within the top 10 % of variable importance projection (VIP) scores (VIP > 1.6). Direction of association with CGE was determined as positive or negative depending on sign of coefficient. “Shared” sub-pathways signifies that gene is part of multiple sub-pathways. CGE, community growth efficiency; AR, aromatic ring; ox., oxidation; ETC, electron transport chain; EPS, extracellular polymeric substance; C, carbon

Within the biosynthesis category, the CGE metric was positively correlated with gene transcript abundance of amino acid biosynthesis (16 genes; p=0.002, r=0.55, Pearson; **Fig. 5a**) and peptidoglycan biosynthesis (3 genes; p=0.03, 0.35, Pearson; **Fig. 5d**), negatively correlated with lipid biosynthesis (3 genes; p=0.002, r=-0.51, Pearson; **Fig. 5b**), and not correlated with nucleotide biosynthesis (7 genes, p=0.3, r=0.17, Pearson; **Fig. 5c**), teichoic acid biosynthesis (2 genes; p=0.2, r=0.29, Pearson; **Fig. S4a**), or EPS biosynthesis (2 genes; p=0.6, r=0.09, Pearson; **Fig. S4b**). Overall, the CGE metric was positively correlated with the sum of gene transcript abundances across all these biosynthesis pathways (total: 33 genes; p=0.004, r=0.50, Pearson; **Fig. 5e**).

**Figure 5.**
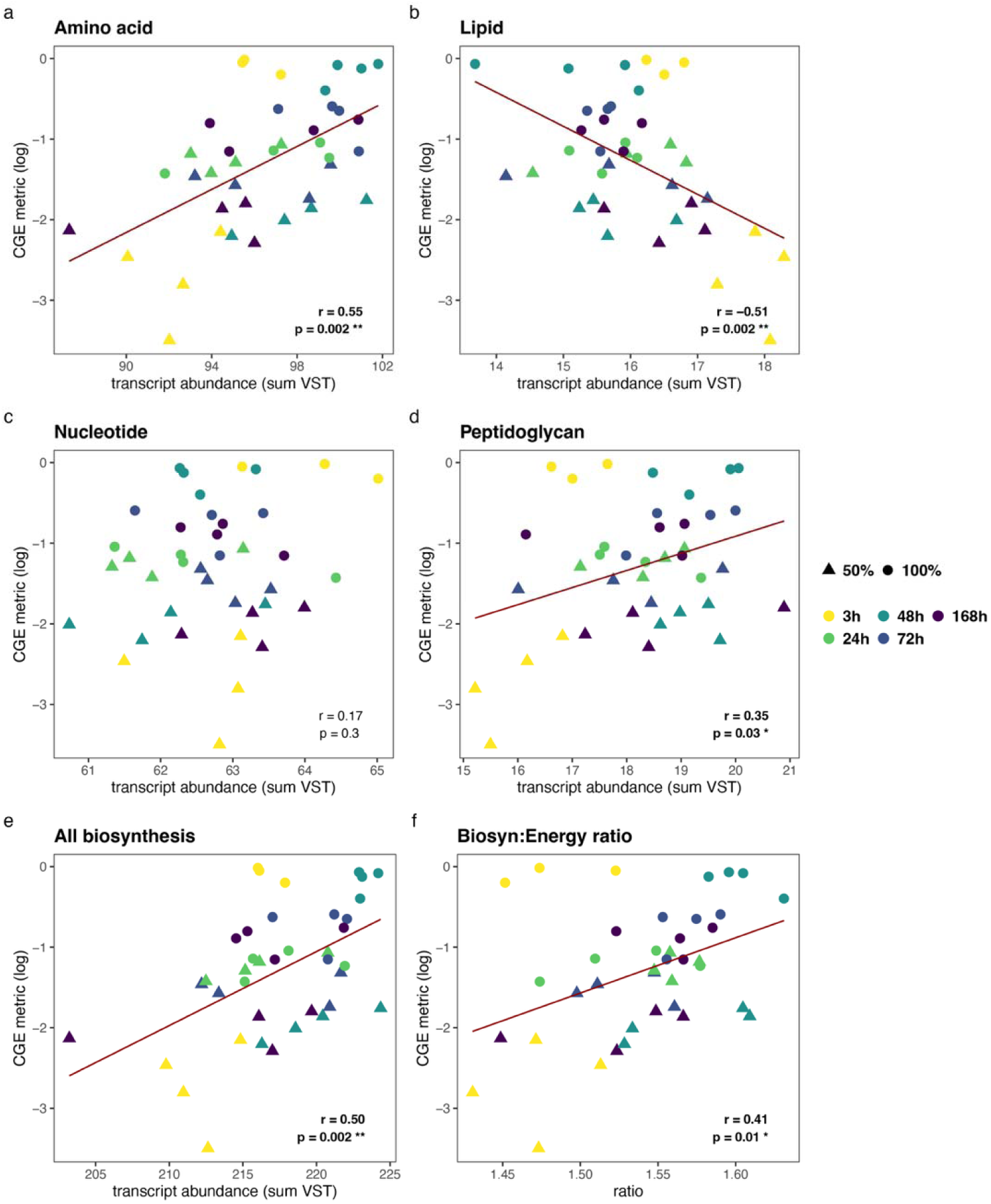
Correlations between community growth efficiency (CGE) metric and transcript abundances (normalized as variance stabilization transformation [VST]) of CGE-associated genes within specific biosynthesis and energy pathways. Biosynthesis pathways include: a) amino acid, b) lipid, c) nucleotide and d) peptidoglycan. e) The sum of transcript abundance across all biosynthesis pathways (amino acid, peptidoglycan, lipids, nucleotides, extracellular polymeric substances (EPS), and teichoic acids. f) Correlation between CGE metric and ratio of transcript abundances of biosynthesis: energy genes. Red line indicates linear regression line for significant correlations. Pearson correlation coefficient (r) and fdr-corrected p-values labeled on each plot. *, p < 0.05; **, p < 0.01.

The CGE metric was not correlated individually with different energy pathways (central C metabolism, fermentation, electron transport chain, or ATPase; p>0.05, **Fig. S4c-f**) nor the sum of transcript abundances across all energy metabolism pathways (total: 19 genes; p=0.6, r=-0.11, Pearson; **Fig. S4h**). There were four genes within the central metabolism pathway associated negatively with CGE (two glycolysis/gluconeogenesis, one pentose phosphate pathway, and one pyruvate oxidation [pyruvate dehydrogenase E1 subunit gene; *PDH*]) compared to one gene associated positively with CGE within the central metabolism pathway involved in the TCA cycle. This suggests that although there were more genes involved in energy metabolism that associated negatively with CGE, the transcript abundances of those positively associated with CGE were greater.

The ratio of gene transcript abundance of biosynthesis to energy production was positively correlated with CGE (**Fig. 5f**), an approximation of the definition of CGE (biosynthesis: energy). A more accurate description would be the sum of the CGE-associated biosynthesis genes to CO_2_-producing genes. The pyruvate dehydrogenase E1 subunit gene (*PDH*; K00163) is the only CO_2_-producing energy gene associated with CGE (negatively) and within the pyruvate oxidation pathway; the ratio of transcript abundance of CGE-associated biosynthesis genes to *PDH* is also positively correlated with CGE (p=0.009, r=0.48, Pearson; **Fig S4i**).

Finally, within the degradation category, the CGE metric was positively correlated with aromatic degradation (28 genes; p = 0.005, r-0.51, Pearson; **Fig. S5d**) and hydrocarbon ring degradation (a subset of aromatic degradation genes that specifically act on rings; 12 genes; p=0.005, r=0.49, Pearson; **Fig. S5e**), negatively correlated with lipid degradation (1 gene [K05939, *aas*], p=0.04, r=-0.35; **Fig. S5b**), and not correlated with amino acid degradation (25 genes; p=0.4, r=0.15, Pearson; **Fig. S5a**) or nucleotide degradation (7 genes; p=0.5, r=-0.08, Pearson; **Fig. S5b**). Overall, the CGE metric was positively correlated with the sum of transcript abundance across all these degradation genes (total: 59 unique genes; p = 0.04, r=0.38, Pearson; **Fig. S5f).**

We screened all CAZy-annotated genes and identified 26 that were strongly associated with CGE, based on VIP scores in the top 10% (ranging from 1.6–3.5; **Fig. S6, Table S3**). Of these, 12 genes were positively associated with CGE, while 14 were negatively associated with CGE. The gene most positively associated with CGE (VIP = 3.5), was a glycosyltransferase (GT12) involved in biosynthesis. Pfam annotation of this gene (PF13641.10; glycosyltransferase like family 2), indicates a role in bacterial capsule formation. Three additional biosynthesis genes (GT76, GT33, and GT22) were also positively associated with CGE and play a role in glycan biosynthesis, a component of cell walls. While glycans are found in microbes and plants, these genes are likely indicative of microbial biosynthesis because plant genes represented < 12.5 % of the sequences (**Fig. S7)** and plants were not growing during the lab wet-up experiment. There was one glycan biosynthesis gene (GT10) that was associated negatively with CGE. The CAZy gene that was most negatively associated with CGE was PL42 (VIP = 2.7), a polysaccharide lyase (rhamnose-a-1,4-D-glucuronate lyase) involved in pectin degradation, a structural component of plant tissues. Most of the remaining CGE-associated genes encoded glycoside hydrolase (GH) enzymes that degrade glycosidic bonds in different substrates to release glucose. Of the GH enzymes associated negatively with CGE, four were specific to plant material substrates (arabinans, xylans, glucuronoxylans), seven to substrates that are found in both plants and microbes (glucans, trehalose, glycans, and galactans), and one that is specific to microbial substrates (chitosan). In contrast, of the GH (and one PL) enzyme associated positively with CGE, three were specific to plant material substrates (arabinans and pectin), two to substrates that are found in both plants and microbes (glycans and galactans), and two that are specific to microbial substrates (mannans and alginate). In addition to the greater number of CAZy genes that were negatively associated with CGE, the total gene transcript abundance (normalized as VST) was also greater (7.3 ± 2.3 vs. 5.3 ± 1.2 [VST-counts]; p = 2.2e-16, LME).

## 4. Discussion

In this study, we found that reduced precipitation during the winter growing season in a Mediterranean grassland ecosystem had lasting effects on SOC composition and microbial growth and mortality dynamics during the subsequent rewetting and Birch effect. These legacy effects persisted even six months later, after the summer dry period, when the soil moisture no longer differed between the treatments. Specifically, reduced legacy precipitation resulted in: 1) Microbial turnover (growth and mortality rates) that was up to two orders of magnitude lower, despite unchanged respiration rates, leading to a decreased community growth efficiency (CGE), and 2) compounds with decreased GFE and increased NOSC. We also identified microbial functions associated with differences in CGEs between legacy precipitation treatments. A lower CGE was linked to decreased biosynthesis, yet no changes in overall energy metabolism, leading to a decreased ratio of transcripts involved in biosynthesis to energy metabolism. These results suggest that reduced precipitation during the growing season—a scenario expected to become more frequent under climate change—not only alters SOC composition but also impacts microbial growth and metabolism during rewetting after the summer dry season. Under reduced legacy precipitation, microbes are less efficient at incorporating carbon into biomass while maintaining similar respiration rates, leading to similar C loss from the soil yet reduced SOC replenishment through microbial turnover.

### Reduced legacy precipitation decreases microbial turnover

Despite similar soil moisture conditions following the summer dry down, we observed that both microbial growth rates and mortality rates were substantially lower—by one and two orders of magnitude, respectively—during rewetting after a single growing season with 50% reduced precipitation. This decrease in microbial growth in the California grassland soil is consistent with previous findings, where growth rates at the drier Sedgwick site (383 mm y^-1^ MAP) were lower than those at the wetter Hopland site (956 mm yr^-1^ MAP), corresponding to a ∼60% decrease in precipitation and shorter C residence times [76]. These results suggest that, over multiple growing seasons with reduced rainfall, the observed decreases in microbial growth and mortality at Hopland would likely lead to reduced soil C accumulation. Given that microbial necromass can contribute up to 50% of SOC in grassland ecosystems [77], a consistent two-order magnitude decrease in mortality each year would significantly reduce necromass inputs to SOC.

The observed changes in microbial growth may be linked to shifts in the composition and physiological state of the microbial community induced by legacy precipitation conditions. Under reduced legacy precipitation, the relative abundance of *Actinobacteria* was greater, that of *Proteobacteria* was lower, and that of *Firmicutes* remained constant. However, *Firmicutes*, which made up only ∼5% of the total community before rewetting, accounted for ∼20% and 40% of the growing community at 3 h post rewetting in the normal and reduced legacy precipitation treatments, respectively. This pattern is consistent with previous findings at this site, where *Firmicutes* were also a major component of the growing community after being rewet [6, 77, 78]. In contrast, *Actinobacteria* made up a greater proportion (∼40%) of the total community under reduced legacy precipitation but only ∼20% of the growing community at 3 h post rewetting. Both *Firmicutes* and *Actinobacteria* are recognized as drought-tolerant taxa [6, 78, 79]. However, their survival strategies likely differ: *Actinobacteria* may persist with some active growth [78], whereas *Firmicutes* typically survive by forming dormant spores [79]. We propose that rewetting triggers the exit from dormancy in *Firmicutes*, leading to rapid growth, whereas *Actinobacteria* may experience mortality due to rewetting stress after a prolonged period of low activity during the dry season. Overall, our findings highlight that legacy precipitation conditions distinctly shape the post-rewetting growth dynamics of *Firmicutes* and *Actinobacteria*, with *Firmicutes* exhibiting enhanced recovery and *Actinobacteria* displaying reduced growth under reduced precipitation regimes.

These changes in taxonomic turnover have implications not only for growth dynamics but also for the biochemical nature of microbial necromass. Under normal legacy precipitation, dying taxa were dominated by gram-negative *Proteobacteria*, whereas under reduced legacy precipitation, gram-positive Actinobacteria dominated. This suggests a shift in necromass composition from gram-negative to gram-positive origins, which differ in their macromolecule compositions (e.g., peptidoglycans, glycoproteins, lipoproteins, and phospholipids). Because cell wall components can adsorb onto mineral surfaces [80] and the strength of this binding varies among different macromolecules, these changes may influence both the type of compounds [81] entering the soil carbon pool and their potential for stabilization.

Importantly, fungi likely contributed to microbial biomass turnover and respiration in this study. While we did not measure fungal growth here, previous work in Hopland soils have shown that fungal and bacterial/archaeal growth respond similarly to rewetting events [79, 81, 82]. Furthermore, transcriptomic analyses indicated comparable ratios of bacteria + archaea to fungi across legacy precipitation treatments and timepoints (**Fig. S7)**, suggesting that fungal growth and mortality likely followed similar patterns across legacy precipitation treatments as those of bacteria and archaea. Future studies that directly quantify fungal growth and turnover will help refine microbial CGE estimates and provide a more complete understanding of how precipitation variability influences oil carbon cycling.

### Influence of bioavailable substrate composition on CGE

Legacy precipitation influenced the composition of the organic carbon pool, altering the types of WEOC substrates available to microbes. Consistent with previous studies, we found that reduced precipitation led to a WEOC pool with fewer unique compounds typical of microbial biomass and necromass, such as lipids, proteins, and amino sugars [80, 82, 83], and more unique compounds derived from plant detritus, including lignin and tannins [84]. This shift to residual plant detritus contributed to an increase in NOSC, indicating that a more oxidized SOC pool was consistent with equal microbial mineralization but reduced new microbial biomass production in the reduced precipitation treatments, resulting in greater C loss as CO_2_.

In addition to substrate GFE, elemental stoichiometry (C:N:P) strongly influences microbial CUE [31, 32]. The stoichiometry of these substrates and their hierarchy of energetic favorability can be approximated as follows: microbial biomass/necromass (global average ∼60:7:1) [86] > plant detritus (grassland belowground ∼200:6.7:1) [19, 86]. Microbial biomass is most favorable because its elemental ratios closely match those of the consuming microbes, minimizing the energetic cost of assimilation. Under normal legacy precipitation conditions, rewetting induces rapid microbial mortality, resulting in the release of a fresh pool of nutrient-rich biomass. This energetically favorable substrate likely contributed to the observed increase in CGE at 3 h post rewetting. In support of this finding, a high CGE coincided with increased expression of genes involved in amino acid, peptidoglycan, and glycolipid biosynthesis, as well as several necromass (bacterial/fungal cell wall)-degrading enzymes (for substrates including mannans and alginate), all of which are indicative of active biomass degradation combined with growth. Previous work at Hopland further supports this mechanism, suggesting that the nutrient-rich products and biomolecules from microbial mortality generated following wet-up can fuel a substantial portion of the microbial response to sudden increases in soil moisture [19, 87]. Furthermore, necromass accrual during the summer dry down could be made bioavailable to the microbes with the sudden increase in moisture; amino sugars, indicative of necromass, were higher before rewetting in the normal compared to the reduced legacy precipitation treatment and decreased significantly by 3 h post rewet indicating rapid microbial consumption.

In contrast, plant detritus imposes a stoichiometric mismatch, requiring microbes to both expend energy-producing extracellular enzymes and respire excess carbon to acquire sufficient nitrogen [32]. In our study, we indeed found that low CGE was associated with greater diversity and expression of genes encoding plant material-degrading enzymes (for substrates including pectin, arabinans, glucuronoxylans, and xylans). This observation aligns with previous studies reporting a negative correlation between enzyme production and CUE [89], suggesting that increased enzyme demand diverts carbon from biomass synthesis to energy generation. Additionally, gene expression data suggests decreased P and N availability under reduced legacy precipitation— conditions that are also associated with lower CUE [32, 88]. In this study, low CGE was specifically associated with the expression of *senX3* [K07768] and *regX3* [K07776], genes activated during phosphate limitation, and *ntrX* [K13599], a gene activated during nitrogen limitation. Together, these findings highlight that legacy precipitation shapes both substrate quality and microbial nutrient stress, ultimately influencing microbial CGE.

### Microbial energy, degradation, and biosynthesis metabolisms associated with CGE

Consistent with our hypothesis that higher CGE would be associated with increased expression of biosynthesis genes, CGE metric increased as transcript abundances for CGE-associated biosynthesis genes in the metatranscriptome increased. However, contrary to our hypothesis that higher CGE would be associated with decreased expression of energy genes, CGE metric did not show an overall correlation with the combined transcript levels of CGE-associated energy metabolism genes. However, as CGE metric increased, the ratio of biosynthesis:energy transcript abundance also increased, indicating decreased transcriptional investment in energy-supplying reactions per unit of anabolic transcription as growth yield increased. At the gene level, more energy genes were negatively associated with CGE metric than positively (13 vs. 8; **Table 1**) and were concentrated in central C metabolism pathways (glycolysis/gluconeogenesis, pentose-phosphate pathway [PPP], and pyruvate oxidation). In contrast, there was one gene in the central C metabolism pathway (TCA cycle) that was positively associated with CGE metric.

Although total biosynthetic activity was higher under normal legacy precipitation, lipid biosynthesis was enriched at low CGE metric, especially for fatty acids, glycerolipids, and glycerophospholipids. Flux through central carbon and PPP pathways can supply both the precursor acetyl-CoA as well as NADPH to power fatty-acid elongation and desaturation [80]. The coordinated increase in lipid biosynthesis and lipid degradation at low CGE metric is most consistent with a maintenance program rather than net biomass production characterized by membrane repair and remodeling and futile lipid cycling under nutrient and legacy-moisture stress. In this context, *fabB* (K00647), a core synthase of the fatty-acid pathway, can support triacylglyceride formation for temporary carbon and energy storage [81] and cell-envelope remodeling to maintain osmotic homeostasis [82]. Moreover, lipid catabolism can return P and C to central metabolism during stress [83] and can be tightly coupled to lipid biosynthesis [84], creating microbial turnover that depresses growth yield. This is characteristic of a futile cycle, where opposing biochemical reactions occur simultaneously resulting in net energy loss without productivity [89].

In contrast, peptidoglycan and amino acid biosynthesis were associated with high CGE metric, consistent with rapid microbial biomass production. A second major characteristic of high CGE was aromatic degradation that funnels carbon directly to the TCA cycle via succinyl-CoA. Entry at succinyl-CoA bypasses glycolysis and pyruvate oxidation, avoiding upstream CO_2_-releasing reactions, and yielding substrate level ATP production. This reaction also yields succinate which can continue through the TCA cycle supplying intermediates for amino acid biosynthesis, particularly oxaloacetate which is upstream from the aspartate-family of amino acids (i.e. alanine, lysine, threonine), the biosynthesis of which was associated with high CGE metric. Amino acids are the building blocks of proteins, which make up 52–68% of the microbial cell composition [90], including peptidoglycan components of the cell wall (specifically alanine and lysine) [91]. Aromatic degradation can also generate intermediates that can act as quorum sensing and interspecies signal molecules that trigger cell-to-cell communication and instigate complex microbial interactions [92]. Here, we found 11 genes that were positively associated with CGE within the Quorum Sensing KEGG pathway (map 2024) [93] as opposed to just two that were negatively associated with CGE metric. We also found genes associated with tryptophan metabolism, another downstream pathway from benzoate degradation, that could produce indole, a microbial signal molecule that can impact both microbial community composition and carbon metabolism. Therefore, in addition to energy and biosynthesis precursors, aromatic degradation could promote microbial communication and aggregate formations.

## Conclusions

Climate change-driven increases in drought and rewetting events have the potential to strongly affect microbial activity and SOC accrual in CA grassland ecosystems. Our study demonstrated that while both normal and reduced winter precipitation conditions produced a pronounced Birch effect—characterized by a CO_2_ pulse following rewetting after the summer dry-down period— reduced precipitation led to markedly lower microbial growth and mortality, resulting in diminished necromass formation. Under typical precipitation conditions, high microbial mortality following a rewet event generates an energetically favorable pool of necromass. This necromass supports subsequent microbial growth which we saw in increased transcript abundances of biosynthesis genes, such as amino acids and peptidoglycan. In contrast, reduced legacy precipitation limits necromass availability and shifts the bioavailable SOC pool toward less energetically favorable substrates, such as decomposed plant detritus. These metabolic adjustments are associated with lower microbial community growth efficiency (**Fig. 6**) and increased investment in stress response, such as lipid degradation coupled with biosynthesis for membrane remodeling. Taken together, these findings highlight that legacy precipitation conditions fundamentally alter microbial turnover, activity, and the composition of SOC. Over time, such changes have the potential to reshape SOC stocks and the carbon balance of grassland soils under future climate scenarios.

**Figure. 6.**
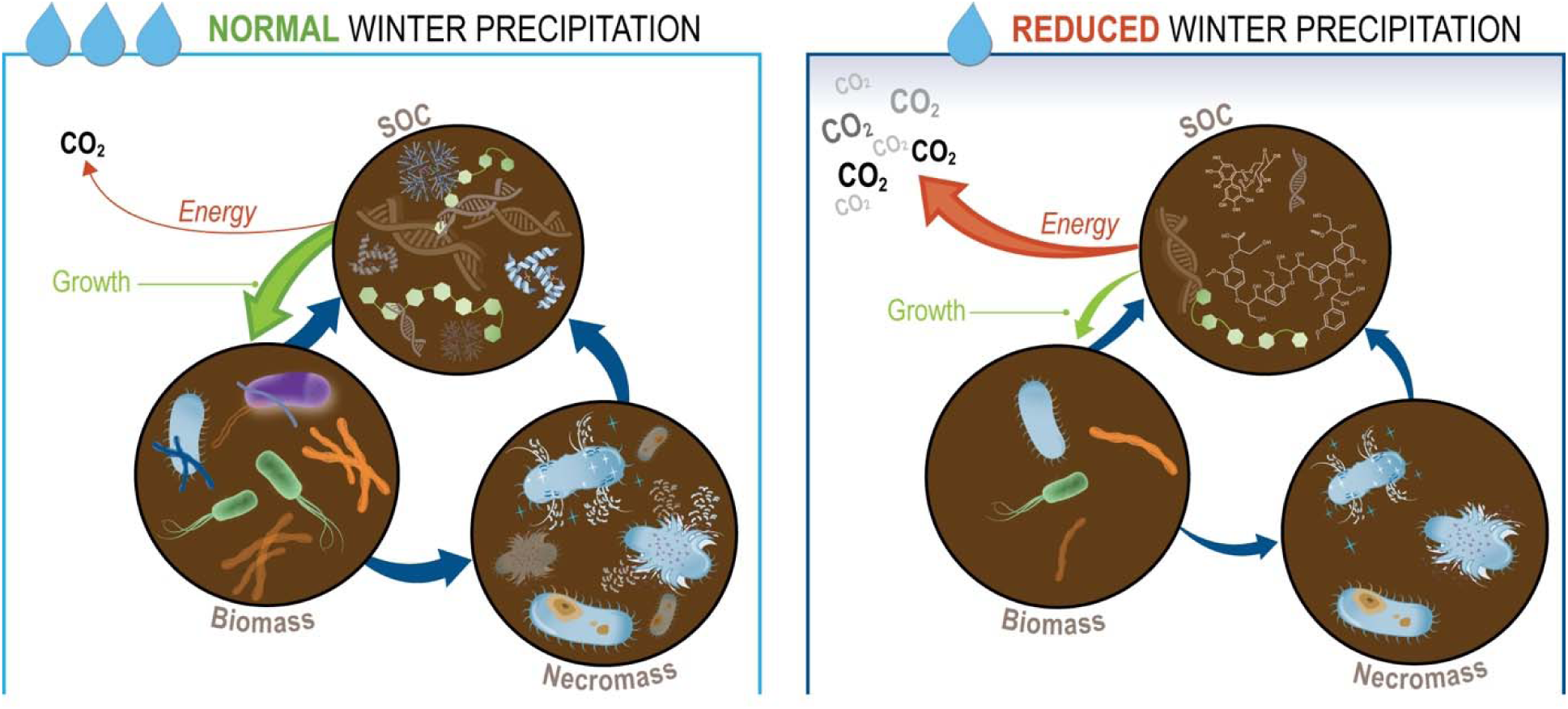
Conceptual diagram showing flow of carbon between soil organic carbon (SOC), biomass, and necromass carbon pools under normal and reduced legacy precipitation. Red lines indicate allocation of carbon to CO_2_ for energy, green lines represents allocation to growth, and blue lines represent general direction and flow of carbon between pools. Thicker lines indicate greater allocation and flow. Molecules drawn within SOC pool represent biomolecules (i.e., DNA, proteins, carbohydrates, lignins, and tannins). Microbial cells are depitcted inside the biomass pool, and cell death is represented in necromass pool. Number of icons in each pool is representative of the quanitity of each in each legacy precipitaiton regime.

## Data and code availability statement

The datasets supporting the conclusions of this article are available as follows: Metatranscriptomics data are available at NCBI Bioproject accession no. PRJNA1088039-PRJNA1088070 and PRJNA1087216-PRJNA1087833 (doi:10.1128/mra.00322-24 [55]). Metagenomics data are available at NCBI Bioproject accession no. PRJNA856348. 16S rRNA gene sequencing data are available at PRJNA1311596. FTICR and CO_2_ data are available at doi:10.6084/m9.figshare.29825273. Code is available at https://github.com/linneakh/CA_grassland_omics.

## Acknowledgements

We thank Jeanette Yuko for creating the conceptual diagram in Figure 6.

## Author contributions

JFB, JPR, MKF, and SJB conceived of and designed the experiment. MMY and KEM helped to run the field study and collect samples. ND and LPT processed samples and collected data for metabolomics. LKH, PFC, JK, and ETS analyzed the data. LKH and SJB wrote the manuscript.

## References

1. Sherwood S, Fu Q. A drier future? Science. 2014;343:737–9. 10.1126/science.1247620.

2. Crowther TW, van den Hoogen J, Wan J, Mayes MA, Keiser AD, Mo L, et al. The global soil community and its influence on biogeochemistry. Science. 2019;365:eaav0550. 10.1126/science.aav0550.

3. Sokol NW, Slessarev E, Marschmann GL, Nicolas A, Blazewicz SJ, Brodie EL, et al. Life and death in the soil microbiome: how ecological processes influence biogeochemistry. Nat Rev Microbiol. 2022. 10.1038/s41579-022-00695-z.

4. Foley MM, Blazewicz SJ, McFarlane KJ, Greenlon A, Hayer M, Kimbrel JA, et al. Active populations and growth of soil microorganisms are framed by mean annual precipitation in three California annual grasslands. Soil Biol Biochem. 2023;177:108886. 10.1016/j.soilbio.2022.108886.

5. Schimel JP. Life in Dry Soils: Effects of Drought on Soil Microbial Communities and Processes. Annu Rev Ecol Evol Syst. 2018;49:409–32. 10.1146/annurev-ecolsys-110617-062614.

6. Honeker LK, Pugliese G, Ingrisch J, Fudyma J, Gil-Loaiza J, Carpenter E, et al. Drought re-routes soil microbial carbon metabolism towards emission of volatile metabolites in an artificial tropical rainforest. Nat Microbiol. 2023;8:1480–94. 10.1038/s41564-023-01432-9.

7. Liang C, Kästner M, Joergensen RG. Microbial necromass on the rise: the growing focus on its role in soil organic matter development. Soil Biol Biochem. 2020;:108000. 10.1016/j.soilbio.2020.108000.

8. Tao F, Huang Y, Hungate BA, Manzoni S, Frey SD, Schmidt MWI, et al. Microbial carbon use efficiency promotes global soil carbon storage. Nature. 2023;618:981–5. 10.1038/s41586-023-06042-3.

9. Pei J, Li J, Luo Y, Rillig MC, Smith P, Gao W, et al. Patterns and drivers of soil microbial carbon use efficiency across soil depths in forest ecosystems. Nat Commun. 2025;16:5218. 10.1038/s41467-025-60594-8.

10. Unlocking Mechanisms for Soil Organic MatterAccumulation: Carbon Use Efficiency and MicrobialNecromass as the Keys.

11. Sokol NW, Foley MM, Blazewicz SJ, Battacharyya A, DiDonato N, Estera-Molina K, et al. The path from root input to mineral-associated soil carbon is shaped by habitat-specific microbial traits and soil moisture. Soil Biol Biochem. 2024;:109367. 10.1016/j.soilbio.2024.109367.

12. He X, Abs E, Allison SD, Tao F, Huang Y, Manzoni S, et al. Emerging multiscale insights on microbial carbon use efficiency in the land carbon cycle. Nat Commun. 2024;15:8010. 10.1038/s41467-024-52160-5.

13. Bouskill NJ, Wood TE, Baran R, Ye Z, Bowen BP, Lim H, et al. Belowground Response to Drought in a Tropical Forest Soil. I. Changes in Microbial Functional Potential and Metabolism. Front Microbiol. 2016;7:525. 10.3389/fmicb.2016.00525.

14. Bouskill NJ, Wood TE, Baran R, Hao Z, Ye Z, Bowen BP, et al. Belowground Response to Drought in a Tropical Forest Soil. II. Change in Microbial Function Impacts Carbon Composition. Front Microbiol. 2016;7:323. 10.3389/fmicb.2016.00323.

15. Malik AA, Swenson T, Weihe C, Morrison EW, Martiny JBH, Brodie EL, et al. Drought and plant litter chemistry alter microbial gene expression and metabolite production. ISME J. 2020;14:2236–47. 10.1038/s41396-020-0683-6.

16. Wu Z, Dijkstra P, Koch GW, Peñuelas J, Hungate BA. Responses of terrestrial ecosystems to temperature and precipitation change: a metaLanalysis of experimental manipulation. Glob Chang Biol. 2011;17:927–42. 10.1111/j.1365-2486.2010.02302.x.

17. Canarini A, Wanek W, Watzka M, Sandén T, Spiegel H, Šantrůček J, et al. Quantifying microbial growth and carbon use efficiency in dry soil environments via 18 O water vapor equilibration. Glob Chang Biol. 2020;26:5333–41. 10.1111/gcb.15168.

18. Simon E, Canarini A, Martin V, Séneca J, Böckle T, Reinthaler D, et al. Microbial growth and carbon use efficiency show seasonal responses in a multifactorial climate change experiment. Commun Biol. 2020;3:584. 10.1038/s42003-020-01317-1.

19. Ullah MR, Carrillo Y, Dijkstra FA. Drought-induced and seasonal variation in carbon use efficiency is associated with fungi:bacteria ratio and enzyme production in a grassland ecosystem. Soil Biol Biochem. 2021;155:108159. 10.1016/j.soilbio.2021.108159.

20. Birch HF. The effect of soil drying on humus decomposition and nitrogen availability. Plant Soil. 1958;10:9–31. 10.1007/bf01343734.

21. Lee X, Wu H-J, Sigler J, Oishi C, Siccama T. Rapid and transient response of soil respiration to rain. Glob Chang Biol. 2004;10:1017–26. 10.1111/j.1529-8817.2003.00787.x.

22. Nguyen NB, Migliavacca M, Bassiouni M, Baldocchi DD, Gherardi LA, Green JK, et al. Widespread underestimation of rain-induced soil carbon emissions from global drylands. Nat Geosci. 2025. 10.1038/s41561-025-01754-9.

23. Blazewicz SJ, Hungate BA, Koch BJ, Nuccio EE, Morrissey E, Brodie EL, et al. Taxon-specific microbial growth and mortality patterns reveal distinct temporal population responses to rewetting in a California grassland soil. ISME J. 2020;14:1520–32. 10.1038/s41396-020-0617-3.

24. Blazewicz SJ, Schwartz E, Firestone MK. Growth and death of bacteria and fungi underlie rainfall-induced carbon dioxide pulses from seasonally dried soil. Ecology. 2014;95:1162–72. 10.1890/13-1031.1.

25. Peterson GD. Contagious disturbance, ecological memory, and the emergence of landscape pattern. Ecosystems. 2002;5:329–38. 10.1007/s10021-001-0077-1.

26. Canarini A, Schmidt H, Fuchslueger L, Martin V, Herbold CW, Zezula D, et al. Ecological memory of recurrent drought modifies soil processes via changes in soil microbial community. Nat Commun. 2021;12:5308. 10.1038/s41467-021-25675-4.

27. Wang B, Allison SD. Drought legacies mediated by trait tradeLoffs in soil microbiomes. Ecosphere. 2021;12. 10.1002/ecs2.3562.

28. Wang S, Thi Thu Hoang D, Anh The Luu, Mostafa T, Razavi BS. Environmental memory of microbes regulates the response of soil enzyme kinetics to extreme water events: Drought-rewetting-flooding. Geoderma. 2023;437:116593. 10.1016/j.geoderma.2023.116593.

29. E. Evans S, D. Allison S, V. Hawkes C. Microbes, memory and moisture: Predicting microbial moisture responses and their impact on carbon cycling. Funct Ecol. 2022;36:1430–41. 10.1111/1365-2435.14034.

30. Liu L, Estiarte M, Bengtson P, Li J, Asensio D, Wallander H, et al. Drought legacies on soil respiration and microbial community in a Mediterranean forest soil under different soil moisture and carbon inputs. Geoderma. 2022;405:115425. 10.1016/j.geoderma.2021.115425.

31. Mendonca CM, Zhang L, Waldbauer JR, Aristilde L. Disproportionate carbon dioxide efflux in bacterial metabolic pathways for different organic substrates leads to variable contribution to carbon-use efficiency. Environ Sci Technol. 2024;58:11041–52. 10.1021/acs.est.4c01328.

32. Manzoni S, Taylor P, Richter A, Porporato A, Ågren GI. Environmental and stoichiometric controls on microbial carbon-use efficiency in soils. New Phytol. 2012;196:79–91. 10.1111/j.1469-8137.2012.04225.x.

33. Geyer KM, Kyker-Snowman E, Grandy AS, Frey SD. Microbial carbon use efficiency: accounting for population, community, and ecosystem-scale controls over the fate of metabolized organic matter. Biogeochemistry. 2016;127:173–88. 10.1007/s10533-016-0191-y.

34. Becchetti T, George M, McDougald N, Flavel D, Vaughn C, Forero L, et al. Rangeland management series: Annual range forage production. University of California, Agriculture and Natural Resources. 2016. 10.3733/ucanr.8018.

35. Fossum C, Estera-Molina KY, Yuan M, Herman DJ, Chu-Jacoby I, Nico PS, et al. Belowground allocation and dynamics of recently fixed plant carbon in a California annual grassland. Soil Biol Biochem. 2022;165:108519. 10.1016/j.soilbio.2021.108519.

36. Sieradzki ET, Greenlon A, Nicolas AM, Firestone MK, Pett-Ridge J, Blazewicz SJ, et al. Functional succesision of actively growing soil microorganisms during rewetting is shaped by precipitation history. Bioarxiv. 2022.

37. Chuckran PF, Estera-Molina K, Nicolas AM, Sieradzki ET, Dijkstra P, Firestone MK, et al. Codon bias, nucleotide selection, and genome size predict in situ bacterial growth rate and transcription in rewetted soil. Proc Natl Acad Sci U S A. 2025;122:e2413032122. 10.1073/pnas.2413032122.

38. Nicolas AM, Sieradzki ET, Pett-Ridge J, Banfield JF, Taga ME, Firestone MK, et al. A subset of viruses thrives following microbial resuscitation during rewetting of a seasonally dry California grassland soil. Nat Commun. 2023;14:5835. 10.1038/s41467-023-40835-4.

39. Barnard RL, Osborne CA, Firestone MK. Changing precipitation pattern alters soil microbial community response to wet-up under a Mediterranean-type climate. ISME J. 2015;9:946–57. 10.1038/ismej.2014.192.

40. Nuccio EE, Blazewicz SJ, Lafler M, Campbell AN, Kakouridis A, Kimbrel JA, et al. HT-SIP: a semi-automated stable isotope probing pipeline identifies cross-kingdom interactions in the hyphosphere of arbuscular mycorrhizal fungi. Microbiome. 2022;10:199. 10.1186/s40168-022-01391-z.

41. Buckley DH, Huangyutitham V, Hsu S-F, Nelson TA. Stable isotope probing with 15N achieved by disentangling the effects of genome G+C content and isotope enrichment on DNA density. Appl Environ Microbiol. 2007;73:3189–95. 10.1128/AEM.02609-06.

42. Fierer N, Jackson JA, Vilgalys R, Jackson RB. Assessment of soil microbial community structure by use of taxon-specific quantitative PCR assays. Appl Environ Microbiol. 2005;71:4117–20. 10.1128/AEM.71.7.4117-4120.2005.

43. Caporaso JG, Lauber CL, Walters WA, Berg-Lyons D, Lozupone CA, Turnbaugh PJ, et al. Global patterns of 16S rRNA diversity at a depth of millions of sequences per sample. Proc Natl Acad Sci U S A. 2011;108 Suppl 1 supplement_1:4516–22. 10.1073/pnas.1000080107.

44. Callahan BJ, McMurdie PJ, Rosen MJ, Han AW, Johnson AJA, Holmes SP. DADA2: High-resolution sample inference from Illumina amplicon data. Nat Methods. 2016;13:581–3. 10.1038/nmeth.3869.

45. Wang Q, Garrity GM, Tiedje JM, Cole JR. Naïve Bayesian classifier for rapid assignment of rRNA sequences into the new bacterial taxonomy. Appl Environ Microbiol. 2007;73:5261–7. 10.1128/aem.00062-07.

46. R core team. R: A language and environment for statistical computing. R Foundation for Statistical Computing, Vienna, Austria; 2024.

47. Hungate BA, Mau RL, Schwartz E, Caporaso JG, Dijkstra P, van Gestel N, et al. Quantitative microbial ecology through stable isotope probing. PeerJ. 2015. 10.7287/peerj.preprints.1282.

48. Koch BJ, McHugh TA, Hayer M, Schwartz E, Blazewicz SJ, Dijkstra P, et al. Estimating taxonLspecific population dynamics in diverse microbial communities. Ecosphere. 2018;9:e02090. 10.1002/ecs2.2090.

49. Martin-Laurent F, Philippot L, Hallet S, Chaussod R, Germon JC, Soulas G, et al. DNA extraction from soils: Old bias for new microbial diversity analysis methods. Appl Environ Microbiol. 2001;67:4397–4397. 10.1128/aem.67.9.4397-4397.2001.

50. Kanagawa T. Bias and artifacts in multitemplate polymerase chain reactions (PCR). J Biosci Bioeng. 2003;96:317–23. 10.1016/S1389-1723(03)90130-7.

51. Kozarewa I, Ning Z, Quail MA, Sanders MJ, Berriman M, Turner DJ. Amplification-free Illumina sequencing-library preparation facilitates improved mapping and assembly of (G+C)-biased genomes. Nat Methods. 2009;6:291–5. 10.1038/nmeth.1311.

52. McMurdie PJ, Holmes S. phyloseq: an R package for reproducible interactive analysis and graphics of microbiome census data. PLoS One. 2013;8:e61217. 10.1371/journal.pone.0061217.

53. Bushnell B, Rood J, Singer E. BBMerge – Accurate paired shotgun read merging via overlap. PLOS ONE. 2017;12:e0185056. 10.1371/journal.pone.0185056.

54. Joshi NA, Fass JN. Sickle: A sliding-window, adaptive, quality-based trimming tool for FastQ files. 2011.

55. Li D, Liu C-M, Luo R, Sadakane K, Lam T-W. MEGAHIT: an ultra-fast single-node solution for large and complex metagenomics assembly via succinct de Bruijn graph. Bioinformatics. 2015;31:1674–6. 10.1093/bioinformatics/btv033.

56. Shaffer M, Borton MA, McGivern BB, Zayed AA, La Rosa SL, Solden LM, et al. DRAM for distilling microbial metabolism to automate the curation of microbiome function. Nucleic Acids Res. 2020;48:8883–900. 10.1093/nar/gkaa621.

57. Aramaki T, Blanc-Mathieu R, Endo H, Ohkubo K, Kanehisa M, Goto S, et al. KofamKOALA: KEGG Ortholog assignment based on profile HMM and adaptive score threshold. Bioinformatics. 2020;36:2251–2. 10.1093/bioinformatics/btz859.

58. Zhang H, Yohe T, Huang L, Entwistle S, Wu P, Yang Z, et al. dbCAN2: a meta server for automated carbohydrate-active enzyme annotation. Nucleic Acids Res. 2018;46:W95–101. 10.1093/nar/gky418.

59. Steinegger M, Söding J. MMseqs2 enables sensitive protein sequence searching for the analysis of massive data sets. Nat Biotechnol. 2017;35:1026–8. 10.1038/nbt.3988.

60. Chuckran PF, Estera-Molina K, Huntemann M, Foster B, Roux S, Mukherjee S, et al. Metatranscriptomes of California grassland soil microbial communities in response to rewetting. Microbiol Resour Announc. 2024;13:e0032224. 10.1128/mra.00322-24.

61. Kopylova E, Noé L, Touzet H. SortMeRNA: fast and accurate filtering of ribosomal RNAs in metatranscriptomic data. Bioinformatics. 2012;28:3211–7. 10.1093/bioinformatics/bts611.

62. Aroney STN, Newell RJP, Nissen JN, Camargo AP, Tyson GW, Woodcroft BJ. CoverM: Read alignment statistics for metagenomics. arXiv [q-bio.GN]. 2025.

63. Wood DE, Lu J, Langmead B. Improved metagenomic analysis with Kraken 2. Genome Biol. 2019;20:257. 10.1186/s13059-019-1891-0.

64. Love MI, Huber W, Anders S. Moderated estimation of fold change and dispersion for RNA-seq data with DESeq2. Genome Biol. 2014;15:550. 10.1186/s13059-014-0550-8.

65. Dittmar T, Koch B, Hertkorn N, Kattner G. A simple and efficient method for the solid-phase extraction of dissolved organic matter (SPE-DOM) from seawater. Limnology and Oceanography: Methods. 2008;6:230–5. 10.4319/lom.2008.6.230.

66. Nicora CD, Burnum-Johnson KE, Nakayasu ES, Casey CP, White RA III, Roy Chowdhury T, et al. The MPLEx protocol for multi-omic analyses of soil samples. J Vis Exp. 2018. 10.3791/57343-v.

67. Folch J, Lees M, Stanley GHS. A simple method for the isolation and purification of total lipides from animal tissues. J Biol Chem. 1957.

68. Tolić N, Liu Y, Liyu A, Shen Y, Tfaily MM, Kujawinski EB, et al. Formularity: software for automated formula assignment of natural and other organic matter from ultrahigh-resolution mass spectra. Anal Chem. 2017;89:12659–65.

69. Ayala-Ortiz C, Graf-Grachet N, Freire-Zapata V, Fudyma J, Hildebrand G, AminiTabrizi R, et al. MetaboDirect: an analytical pipeline for the processing of FT-ICR MS-based metabolomic data. Microbiome. 2023;11:28. 10.1186/s40168-023-01476-3.

70. Tfaily MM, Chu RK, Toyoda J, Tolić N, Robinson EW, Paša-Tolić L, et al. Sequential extraction protocol for organic matter from soils and sediments using high resolution mass spectrometry. Anal Chim Acta. 2017;972:54–61. 10.1016/j.aca.2017.03.031.

71. Kuznetsova A, Brockhoff PB, Christensen RHB. LmerTest package: Tests in linear mixed effects models. J Stat Softw. 2017;82. 10.18637/jss.v082.i13.

72. Lenth RV. emmeans: Estimated Marginal Means, aka Least-Squares Means. R package. 2025.

73. Signorell A. DescTools: Tools for Descriptive Statistics. 2025.

74. Dixon P. VEGAN, a package of R functions for community ecology. J Veg Sci. 2003;14:927–30. 10.1111/j.1654-1103.2003.tb02228.x.

75. Young R, Avneri-Katz S, McKenna A, Chen H, Bahureksa W, Polubesova T, et al. Composition-dependent sorptive fractionation of anthropogenic dissolved organic matter by Fe(III)-montmorillonite. Soil Syst. 2018;2:14. 10.3390/soilsystems2010014.

76. Bahureksa W, Tfaily MM, Boiteau RM, Young RB, Logan MN, McKenna AM, et al. Soil organic matter characterization by Fourier transform ion cyclotron resonance mass spectrometry (FTICR MS): A critical review of sample preparation, analysis, and data interpretation. Environ Sci Technol. 2021;55:9637–56. 10.1021/acs.est.1c01135.

77. Wang B, An S, Liang C, Liu Y, Kuzyakov Y. Microbial necromass as the source of soil organic carbon in global ecosystems. Soil Biol Biochem. 2021;162:108422. 10.1016/j.soilbio.2021.108422.

78. Metze D, Schnecker J, Canarini A, Fuchslueger L, Koch BJ, Stone BW, et al. Microbial growth under drought is confined to distinct taxa and modified by potential future climate conditions. Nat Commun. 2023;14:5895. 10.1038/s41467-023-41524-y.

79. Veach AM, Zeglin LH. Historical drought affects microbial population dynamics and activity during soil drying and re-wet. Microb Ecol. 2020;79:662–74. 10.1007/s00248-019-01432-5.

80. Miltner A, Bombach P, Schmidt-Brücken B, Kästner M. SOM genesis: microbial biomass as a significant source. Biogeochemistry. 2012;111:41–55. 10.1007/s10533-011-9658-z.

81. Pouran HM. Bacterial cell-mineral interface, its impacts on biofilm formation and bioremediation. In: Handbook of Environmental Materials Management. Cham: Springer International Publishing; 2019. p. 535–56. 10.1007/978-3-319-73645-7_80.

82. Wang Q, Ding J, Zhang Z, Liang C, Lambers H, Zhu B, et al. Rhizosphere as a hotspot for microbial necromass deposition into the soil carbon pool. Journal of Ecology. 2025;113:168–79. 10.1111/1365-2745.14448.

83. Glaser B, Turrión M-B, Alef K. Amino sugars and muramic acid—biomarkers for soil microbial community structure analysis. Soil Biol Biochem. 2004;36:399–407. 10.1016/j.soilbio.2003.10.013.

84. Kögel-Knabner I. The macromolecular organic composition of plant and microbial residues as inputs to soil organic matter. Soil Biol Biochem. 2002;34:139–62. 10.1016/s0038-0717(01)00158-4.

85. Cleveland CC, Liptzin D. C:N:P stoichiometry in soil: is there a “Redfield ratio” for the microbial biomass? Biogeochemistry. 2007;85:235–52. 10.1007/s10533-007-9132-0.

86. Heyburn J, McKenzie P, Crawley MJ, Fornara DA. Effects of grassland management on plant C:N:P stoichiometry: implications for soil element cycling and storage. Ecosphere. 2017;8:e01963. 10.1002/ecs2.1963.

87. Mganga KZ, Sietiö O-M, Meyer N, Poeplau C, Adamczyk S, Biasi C, et al. Microbial carbon use efficiency along an altitudinal gradient. Soil Biol Biochem. 2022;173:108799. 10.1016/j.soilbio.2022.108799.

88. Xu Q, Li L, Guo J, Guo H, Liu M, Guo S, et al. Active microbial population dynamics and life strategies drive the enhanced carbon use efficiency in high-organic matter soils. MBio. 2024;15:e0017724. 10.1128/mbio.00177-24.

89. Sharma AK, Khandelwal R, Wolfrum C. Futile cycles: Emerging utility from apparent futility. Cell Metab. 2024;36:1184–203. 10.1016/j.cmet.2024.03.008.

90. Bremer H, Dennis PP. Modulation of chemical composition and other parameters of the cell at different exponential growth rates. EcoSal Plus. 2008;3. 10.1128/ecosal.5.2.3.

91. Pellon G. Biosynthesis of peptidoglycans. Biochimie. 1978;60:XI–XII.

92. Díaz E, Jiménez JI, Nogales J. Aerobic degradation of aromatic compounds. Curr Opin Biotechnol. 2013;24:431–42. 10.1016/j.copbio.2012.10.010.

93. Kanehisa Laboratories. KEGG pathway database. KEGG pathway databse. www.genome.jp/kegg/pathway.html. Accessed 1 Jan 2023.

