## Supplementary figures and images for "Reduced legacy precipitation decreases microbial community growth efficiency and alters soil organic carbon in a California grassland"

### Supplemental Figure 1

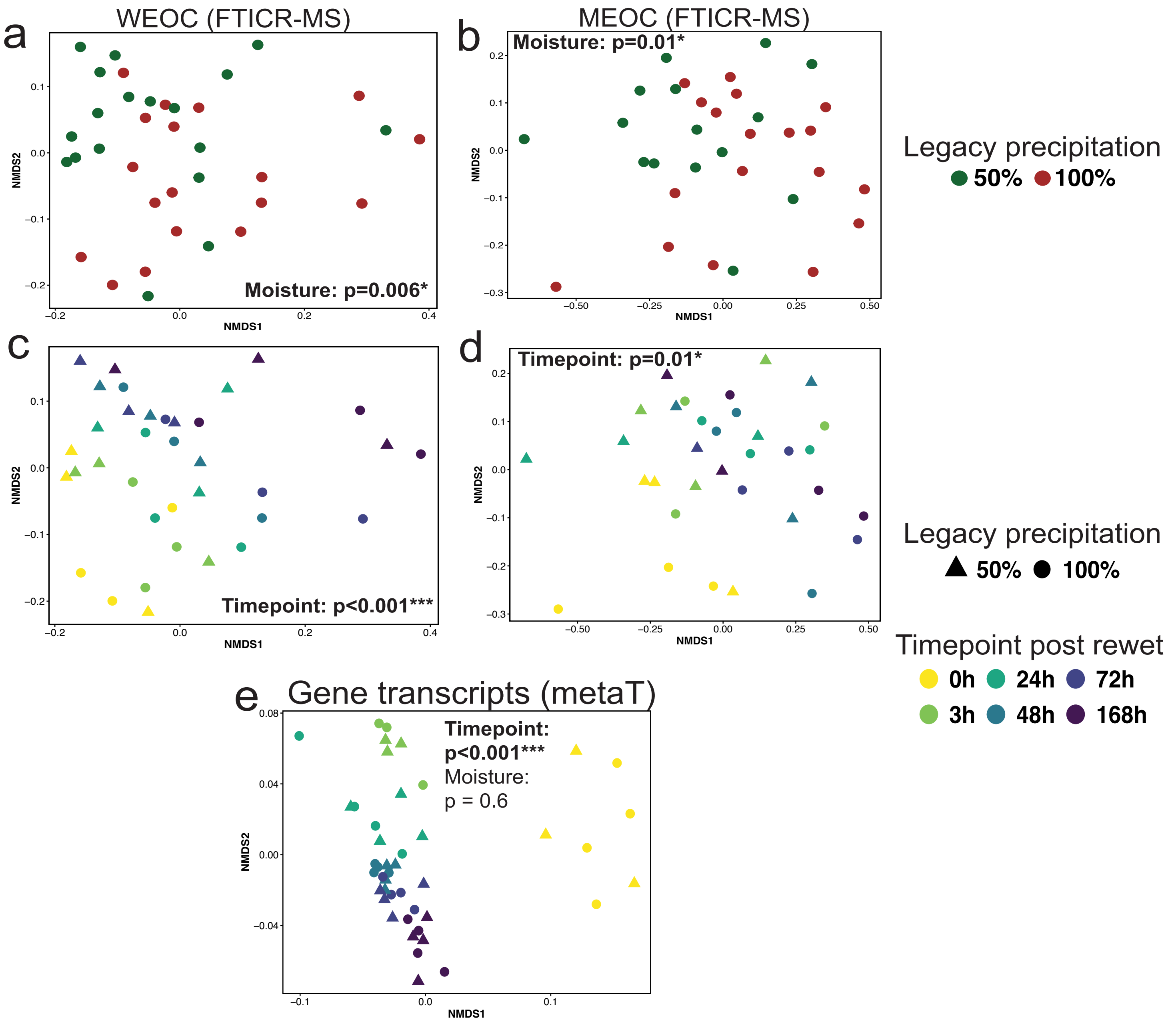

### Supplemental Figure 2

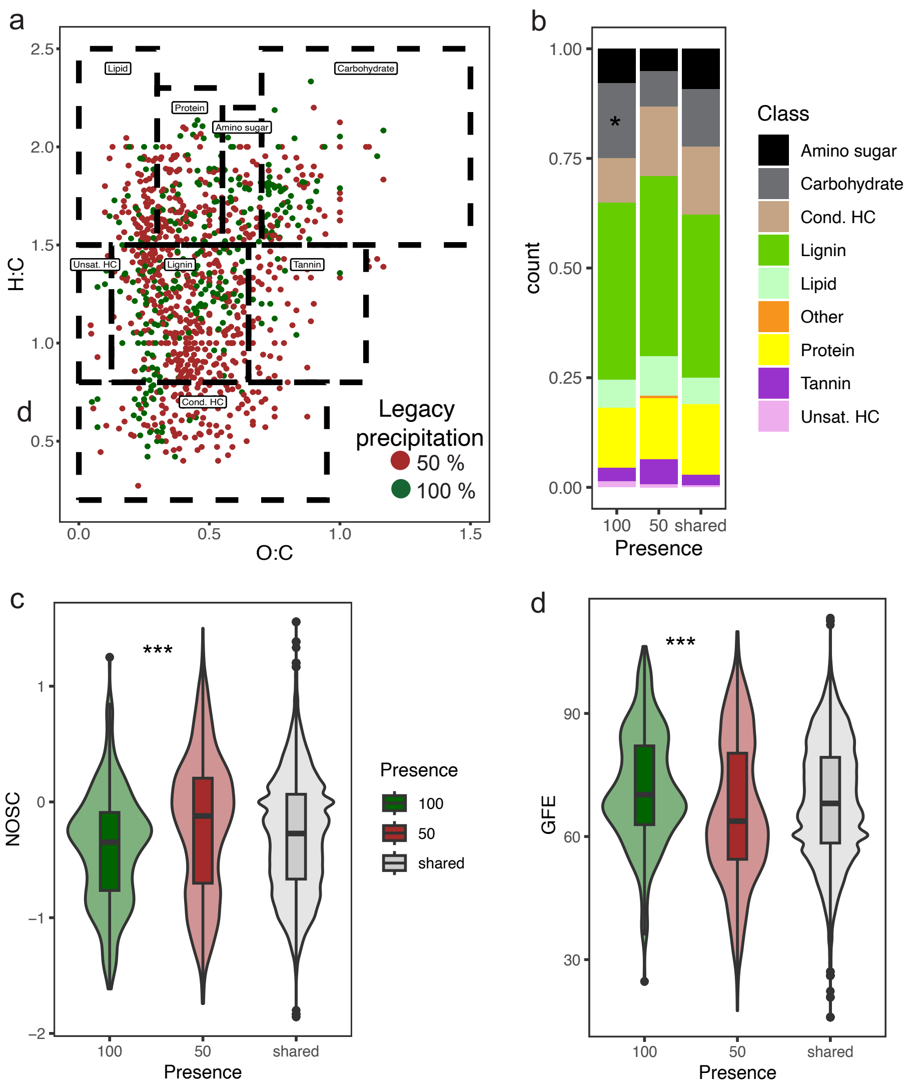

### Supplemental Figure 3

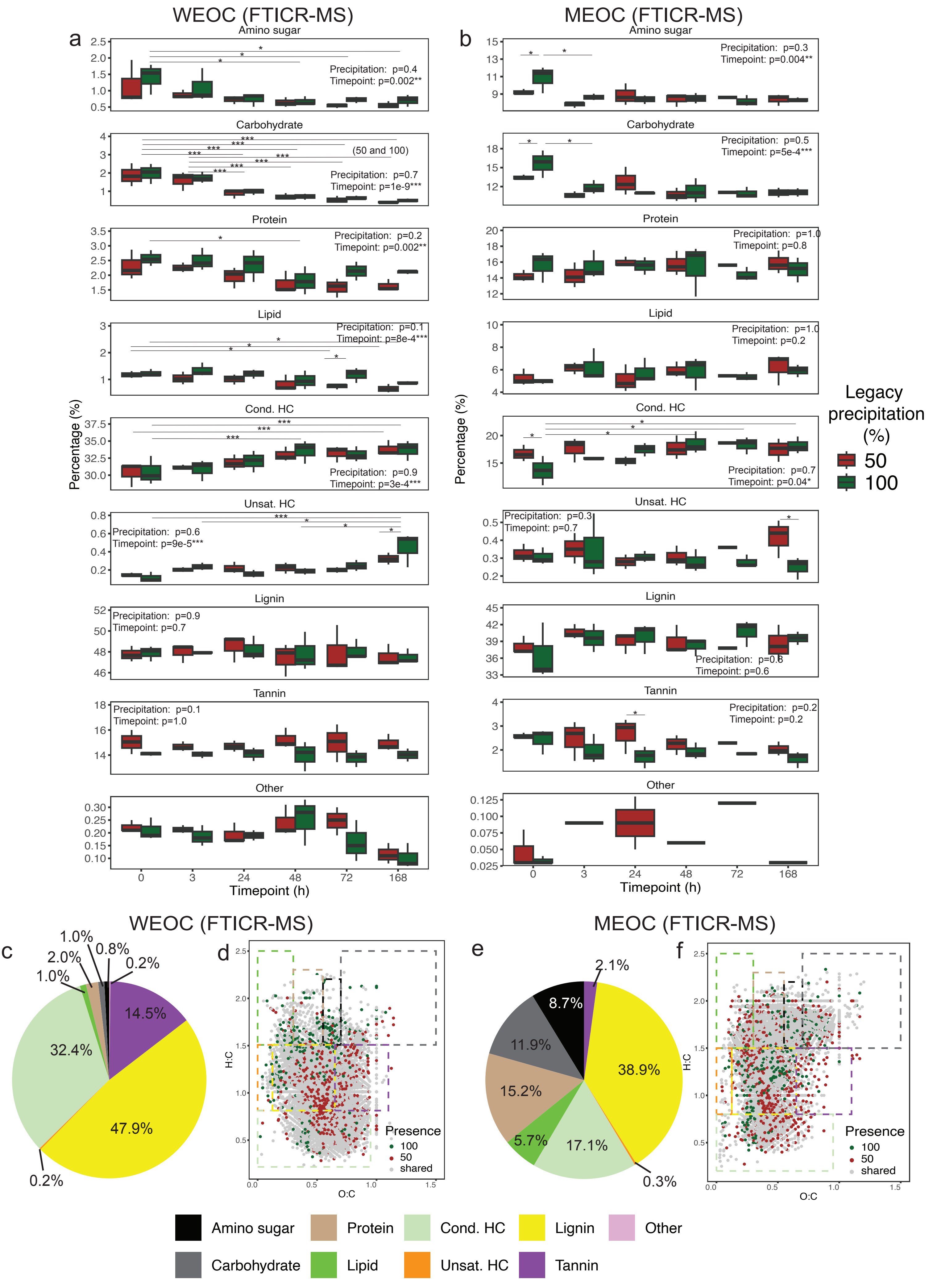

### Supplemental Figure 4

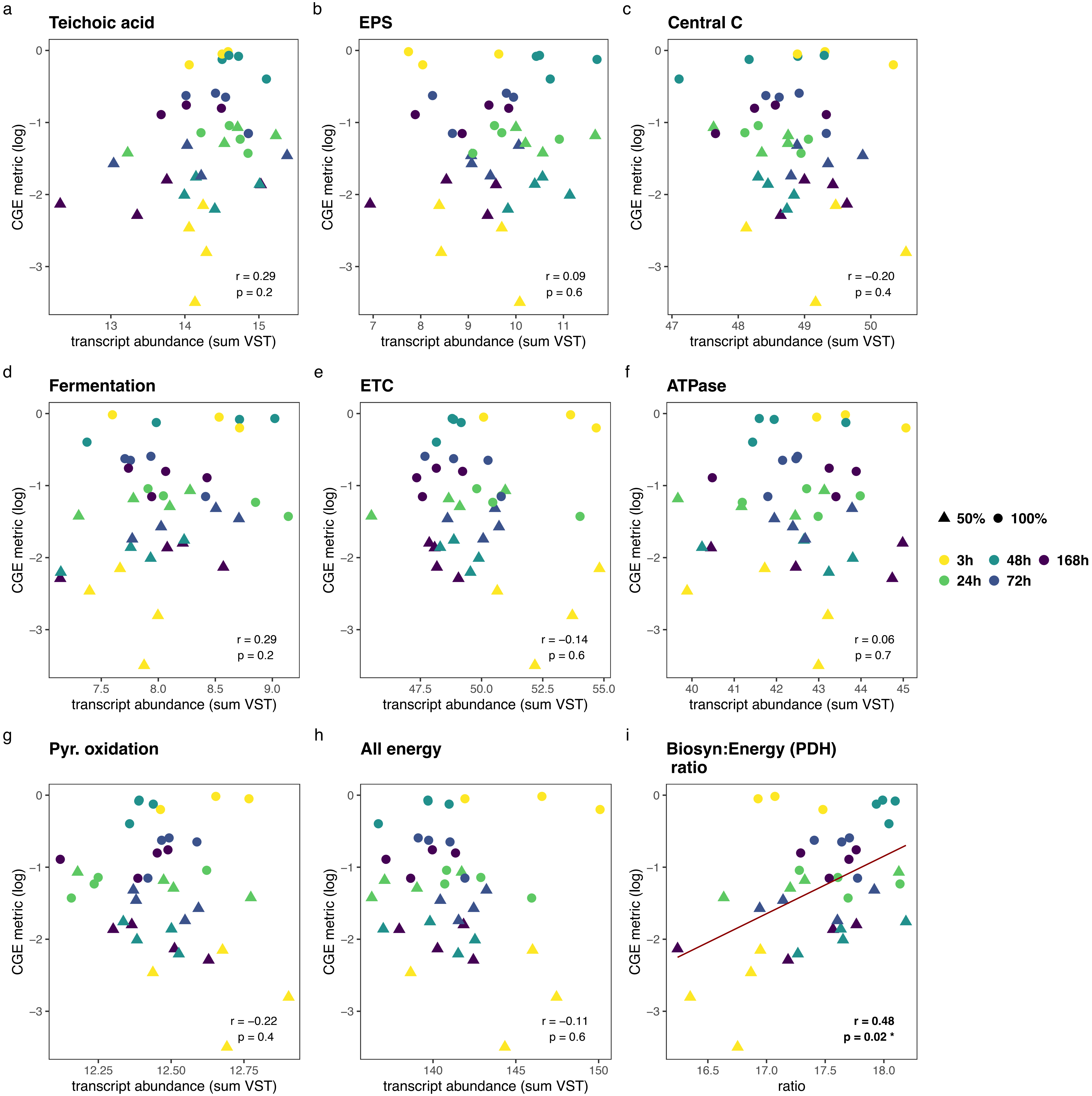

### Supplemental Figure 5

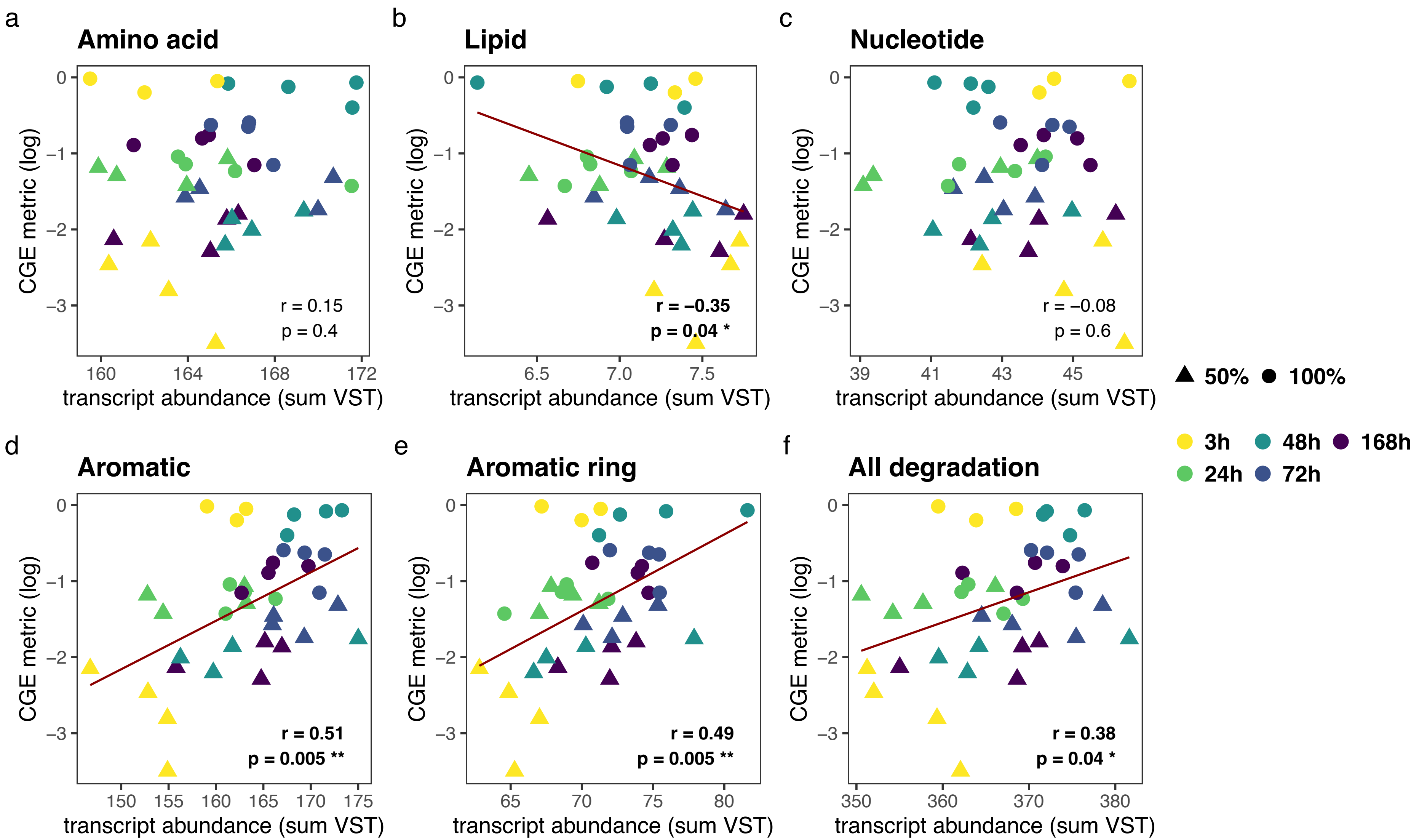

### Supplemental Figure 6

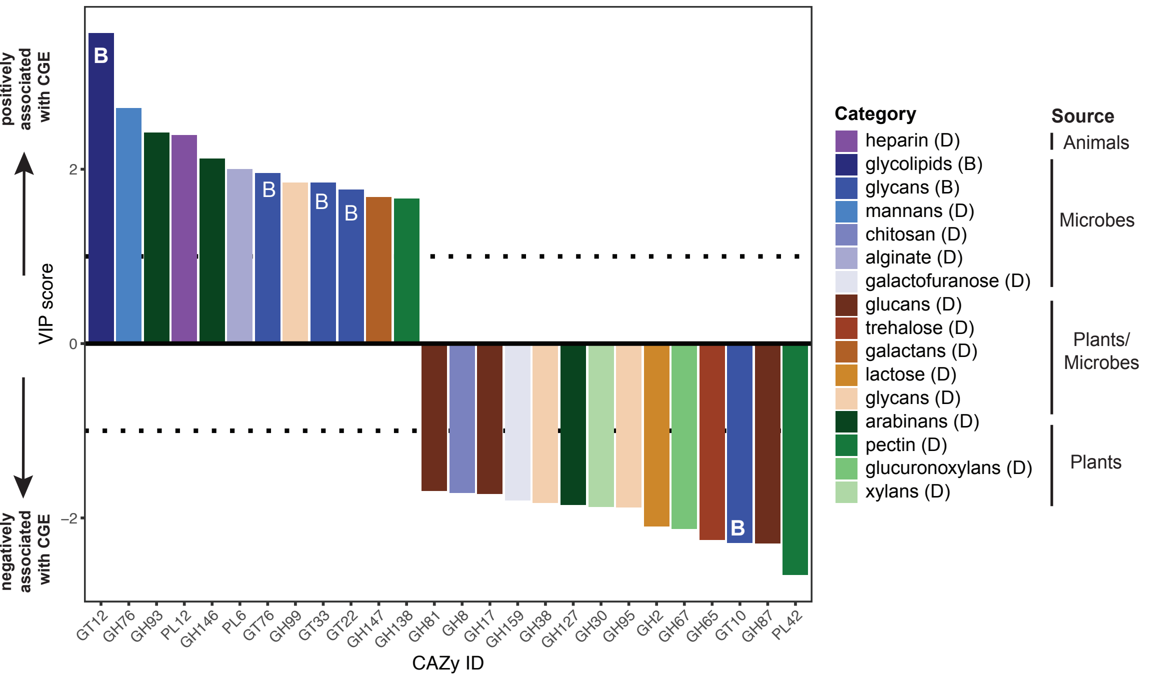

### Supplemental Figure 7

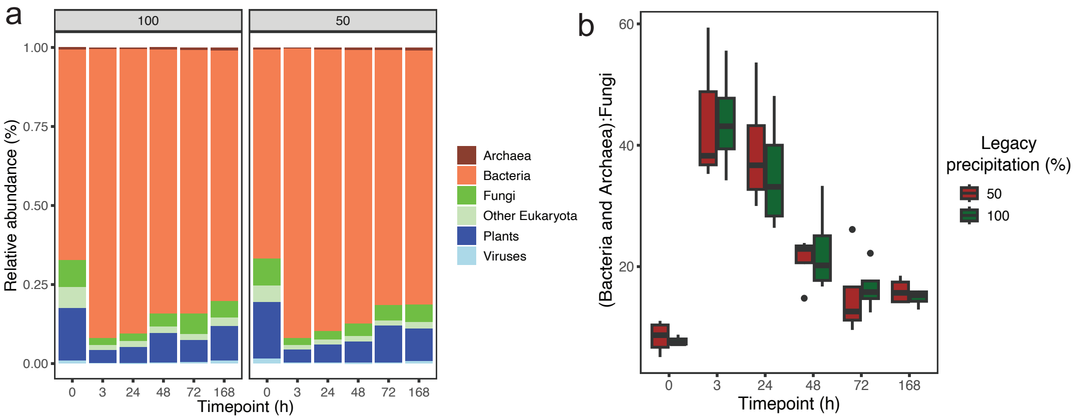
